# A co-operative knock-on mechanism underpins Ca^2+^-selective cation permeation in TRPV channels

**DOI:** 10.1101/2022.04.01.486690

**Authors:** Callum M. Ives, Neil J. Thomson, Ulrich Zachariae

**Affiliations:** Division of Computational Biology, School of Life Sciences, University of Dundee, Dow Street, Dundee, DD1 5EH, UK

## Abstract

The selective exchange of ions across cellular membranes is a vital biological process. Ca^2+^-mediated signalling is implicated in a broad array of physiological processes in cells, whilst elevated intracellular concentrations of Ca^2+^ are cytotoxic. Due to the significance of this cation, strict Ca^2+^ concentration gradients are maintained across the plasma and organelle membranes. Therefore, Ca^2+^ signalling relies on permeation through selective ion channels that control the flux of Ca^2+^ ions. A key family of Ca^2+^-permeable membrane channels are the polymodal signal-detecting Transient Receptor Potential (TRP) ion channels. TRP channels are activated by a wide variety of cues including temperature, small molecules, transmembrane voltage and mechanical stimuli. Whilst most members of this family permeate a broad range of cations non-selectively, TRPV5 and TRPV6 are unique due to their strong Ca^2+^-selectivity. Here, we address the question of how some members of the TRPV subfamily show a high degree of Ca^2+^-selectivity whilst others conduct a wider spectrum of cations. We present results from all-atom molecular dynamics simulations of ion permeation through two Ca^2+^-selective and two non-selective TRPV channels. Using a new method to quantify permeation co-operativity based on mutual information, we show that Ca^2+^-selective TRPV channel permeation occurs by a three binding site knock-on mechanism, whereas a two binding site knock-on mechanism is observed in non-selective TRPV channels. Each of the ion binding sites involved displayed greater affinity for Ca^2+^ over Na^+^. As such, our results suggest that coupling to an extra binding site in the Ca^2+^-selective TRPV channels underpins their increased selectivity for Ca^2+^ over Na^+^ ions. Furthermore, analysis of all available TRPV channel structures shows that the selectivity filter entrance region is wider for the non-selective TRPV channels, slightly destabilising ion binding at this site, which is likely to underlie mechanistic decoupling.

## Introduction

The significance of Ca^2+^ in cellular function was first recognised by Sydney Ringer in 1883, who demonstrated that minute amounts of calcium were required for the contraction of cardiac muscle (1). Ca^2+^ is now recognised as a versatile signalling agent, with cellular Ca^2+^ concentrations impacting a broad array of physiological processes ranging from cell proliferation to cell suicide (2–5). However, the cytoplasmic concentration of Ca^2+^ ions is usually kept low due to cytotoxic consequences (6). Therefore, the controlled opening of channels in cellular and organellar membranes is one of the required mechanisms to allow the influx of this ion from the exoplasm and internal storage compartments into the cytoplasm; this subsequently initiates the Ca^2+^ signalling cascade. The question of how Ca^2+^ channels selectively permeate Ca^2+^ in low concentrations over vastly more abundant Na^+^ ions, and yet conduct them at high rates, has been a longstanding matter of fascination for ion channel researchers (7).

A key example of ion channels that mediate Ca^2+^ permeation across the cytoplasmic membrane is the transient receptor potential (TRP) channel superfamily. In their open state, these polymodal signal-detecting TRP channels allow the transmembrane flux of cations down their electrochemical gradient, thereby increasing the intracellular Ca^2+^ and Na^+^ concentration (8). The malfunction of TRP channels underlies a wide range of pathologies, and they are therefore of immense biomedical importance, serving as drug targets for a variety of existing and candidate drugs (9).

TRP channels assemble primarily as homotetramers to form functional ion channels. A conserved structural feature across all TRP channels is the presence of six transmembrane helices (S1-S6) per subunit, forming two distinct transmembrane domains; a four-helix bundle comprising of helices S1-S4 forming the voltage-sensing like domain (VSLD), and the pore-forming domain consisting of helices S5 and S6 (10). A four-residue ion selectivity filter (SF) is located at the entrance of the channel pore (Figure 1). In addition to this conserved transmembrane architecture, members of the TRP superfamily display highly diverse extramembrane loops and *N*- and *C*-terminal domains between the different subfamilies (11).

**Fig. 1.**
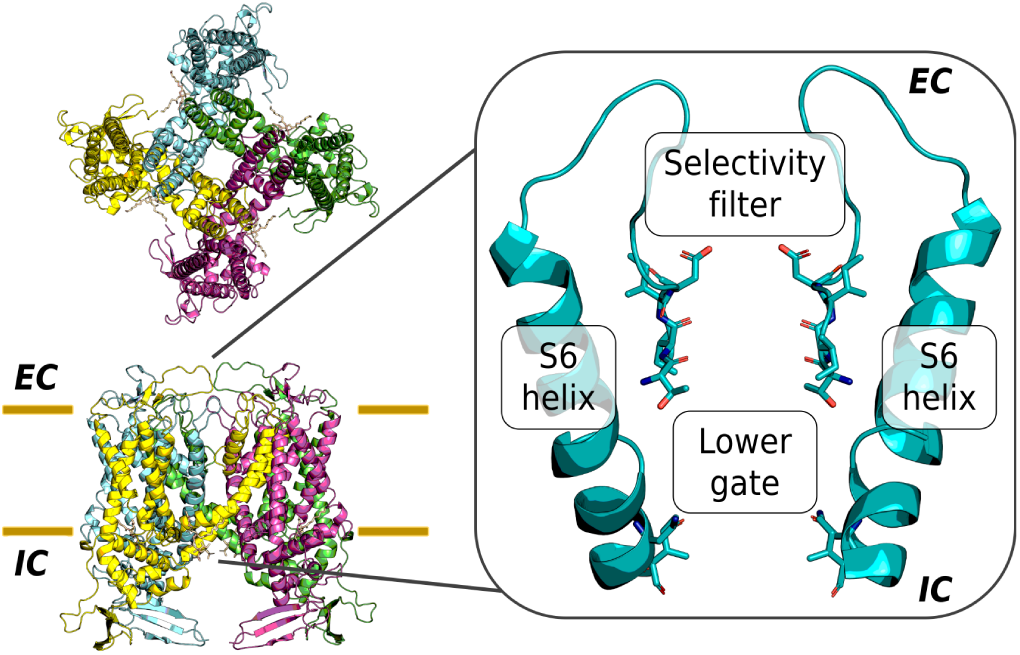
Structure of the truncated construct of TRPV5 of *Oryctolagus cuniculus* used within this work, from the extracellular side (*top-left*) and in the plane of the lipid bilayer (*bottom-left*). In this study, the pore is defined as the region between the constrictions of the channel, namely the top residue of the SF (referred to as the *α*-position of the SF) and the hydrophobic lower gate (*right*).

TRP channels are gated open by a particularly wide range of stimuli, which include temperature, small molecules, transmembrane voltage changes, and mechanical cues (12–14). This superfamily of genes can be divided into seven main subfamilies: TRPA (ankyrin), TRPC (canonical), TRPM (melastatin), TRPML (mucolipin), TRPN (no mechanoreceptor potential C), TRPP (polycystin), and TRPV (vanilloid). It should be noted however, that several less-well characterised TRP subfamilies have also been reported, including the TRPY (15–17), TRPVL (18), and TRPS (19) subfamilies. The TRPV channels are the most intensely studied channel subfamily.

Whilst they are all cation-selective, most TRP channels electrophysiologically characterised to date show only limited discrimination between cation types, as well as between divalent and monovalent cations. However, the TRPV5 and TRPV6 channels are unique due to their high selectivity for Ca^2+^ cations over Na^+^ cations (*P*_*Ca*_*/P*_*Na*_ ∼100:1 from reversal potential measurements) (20, 21). Phylogenetic analysis has demonstrated that the TRPV5 and TRPV6 channels of vertebrates originated from an ancestral TRPV5/6 gene, which then diverged to form TRPV5 and TRPV6 from a duplication event after speciation (22). Both of these channels are constitutively active due to basal levels of phosphatidylinositol 4,5-bisphosphate (PI(4,5)P_2_) in the cellular membrane, and play a key role in Ca^2+^ homeostasis in the body (23). Despite their characteristic Ca^2+^-selectivity, both channels have been shown to permeate monovalent cations such as Na^+^ when divalent cations are absent (20, 24–27).

By contrast, the remaining members of the TRPV subfamily, TRPV1-4, permeate both Ca^2+^ and Na^+^ cations, even in the presence of high Ca^2+^ concentrations, although they are still slightly Ca^2+^-selective, with a permeability ratio *P*_*Ca*_*/P*_*Na*_ ∼ 10:1 (28). These channels gate in response to a number of stimuli, including raised temperature – in particular the archetypal member TRPV1 (12–14), which has led to TRPV1-4 being referred to as thermoTRPV channels – as well as endogenous and exogenous ligands.

In recent years, MD simulations have been successfully employed to shed light on ion channel function and the mode of action of channel-acting drugs in atomistic detail, for instance on K^+^ channels (29, 30), Na^+^ channels (31, 32), Cl^-^ channels (33, 34), and ligand-gated ion channels (35). However, in the case of Ca^2+^-permeating channels, the conventional point-charge models used to describe uncoordinated Ca^2+^ ions have historically been inaccurate due to the neglected effects of electronic polarisation (36). This has resulted in an over-estimation of the binding energies between Ca^2+^ and proteins, hindering an accurate study of Ca^2+^ permeation in channels. Previous efforts to resolve this overestimation have included polarisable force field approaches (37) and re-scaled Ca^2+^ charges (36, 38). More recently, Zhang *et al*. published a new multi-site Ca^2+^ model specifically optimised for Ca^2+^-protein interactions (39), which also showed improved interactions with water. The new model has been employed to characterise the Ca^2+^ permeation and selectivity mechanisms of the type-1 ryanodine receptor (RyR1), accurately reproducing the experimentally recorded conductance of the channel (39, 40).

In the present work, we set out to elucidate the molecular basis of Ca^2+^-selectivity and permeation in the TRPV channel subfamily. We conducted atomistic molecular dynamics (MD) simulations of TRPV channels under transmembrane voltage, and compared the cation permeation mechanism observed in the Ca^2+^-selective TRPV5 and TRPV6 channels to the permeation mechanism in two exemplar non-selective TRPV channels, TRPV2 and TRPV3. In total, we observed 2,849 full ion traversals from 16.95 *µ*s of MD simulations, allowing us to decipher the permeation mechanisms and principles of ion selectivity in the TRPV family with statistical power. Our findings suggest that ion conduction in TRPV channels proceeds via a co-operative knock-on mechanism involving multiple ion binding sites. The degree of co-operativity in ion permeation, linking the multiple binding sites, determines the degree of ion selectivity in the channels.

## Results

### Continuous permeation of Ca^2+^ and Na^+^ in open-state TRPV5 and TRPV6 channels

We performed MD simulations of the pore domain of open-state TRPV5 (41) and TRPV6 (42) channels embedded in POPC (1-palmitoyl-2-oleoyl-sn-glycerol-3-phosphocholine) lipid bilayers under transmembrane voltage (∼ -410 mV). The aqueous solutions contained either 150 mM CaCl_2_ or 150 mM NaCl (herein referred to as mono-cationic solutions), or a mixture consisting of 75 mM CaCl_2_ and 75 mM NaCl (herein referred to as di-cationic solutions). All simulations performed with Ca^2+^ ions utilised the multi-site Ca^2+^ model developed by Zhang *et al*. (39), unless otherwise stated. In both the mono-cationic and the di-cationic solutions, the applied voltage drove a continuous flow of permeating ions through all investigated open-state TRPV channels. Overall, we recorded 433 complete inward channel crossings for Ca^2+^ and 417 for Na^+^ in simulations of the Ca^2+^-selective TRPV channels (Table 1).

**Table 1.** Calculated conductances from MD simulations of ion permeation in Ca^2+^-selective TRPV channels. Mean inward conductances and standard error of the mean (SEM) were calculated from overlapping 50 ns windows from five-fold replicated 250 ns simulations. Permeation of mono-cationic Ca^2+^ or Na^+^ cations was simulated in a solution of 150 mM CaCl_2_ or 150 mM NaCl, respectively. Permeation of a di-cationic mixture of Ca^2+^ and Na^+^ cations was investigated in a solution of 75 mM CaCl_2_ and 75 mM NaCl.

|  | Conductance (pS) |  |  |  |
| --- | --- | --- | --- | --- |
| | $\text{Ca}^{2+}$ | $\text{Na}^+$ | $\text{Ca}^{2+}$ & $\text{Na}^+$ | Literature |
| TRPV5 | $53 \pm 7$ | $49 \pm 6$ | $92 \pm 10$ | $59 \pm 6$ ($\text{Na}^+$) (43) |
| TRPV6 | $117 \pm 12$ | $61 \pm 6$ | $29 \pm 6$ | $58 \pm 4$ ($\text{Na}^+$) (21) |

In TRPV5 and TRPV6, Ca^2+^ ions traversed the entire pore with average time spans of 28.4 ± 3.9 ns (TRPV5) and 12.0 ± 1.0 ns (TRPV6) (Table S4). The calculated Ca^2+^ and Na^+^ conductances from our simulations are shown in (Table 1). The considerable Na^+^ conductances we observed agree with the experimental finding that the highly Ca^2+^-selective TRPV channels conduct Na^+^ well in the absence of Ca^2+^ (20, 24–27). Notably, these conductances are in quantitative agreement with published values measured for Na^+^ *in vitro* (21, 43).

By contrast, control simulations of TRPV5 using the default CHARMM36m force field parameters for Ca^2+^, but otherwise identical conditions, did not exhibit ion permeation; instead, the Ca^2+^ ions remained tightly bound to the protein ion binding sites for the entire course of the simulations. This observation is reflective of the shortcomings of standard parameters for divalent cations in fixed-point charge force fields and highlights the improved accuracy of multi-site Ca^2+^ models in simulating divalent cation permeation and reproducing *in vitro* conductances. In addition, no Cl^-^ anions were observed to permeate TRPV channels in any of our simulations.

### Pore cation binding sites and their preference for Ca^2+^-binding

Prior to the determination of the atomic structures of Ca^2+^ channels and the development of channel-permeable models for Ca^2+^ ions, it had been suggested from experimental observations that Ca^2+^ channels may obtain their selectivity through competition, *i*.*e*. by divalent cations, such as Ca^2+^, binding more strongly at their ion binding sites than monovalent cations, such as Na^+^ (7).

By analysing the individual traces of permeating Ca^2+^ cations along the pore axis *z* of TRPV5 and TRPV6 over time, we identified three cation binding sites inside the channels (Figure 2). We refer to these cation binding sites as sites A, B and C, viewed from the extracellular entrance of the channel selectivity filter to the hydrophobic lower gate. The three cation binding sites were further confirmed by 3D density analysis using PENSA (44) (Figure S2). The PENSA analysis also identified a number of cation binding sites outside of the pore within the extracellular loops of both TRPV5 and TRPV6 (Figure S2), in line with the previous suggestion that TRPV6 contains negatively charged “recruitment sites” that funnel cations towards the entrance of the pore (45, 46).

**Fig. 2.**
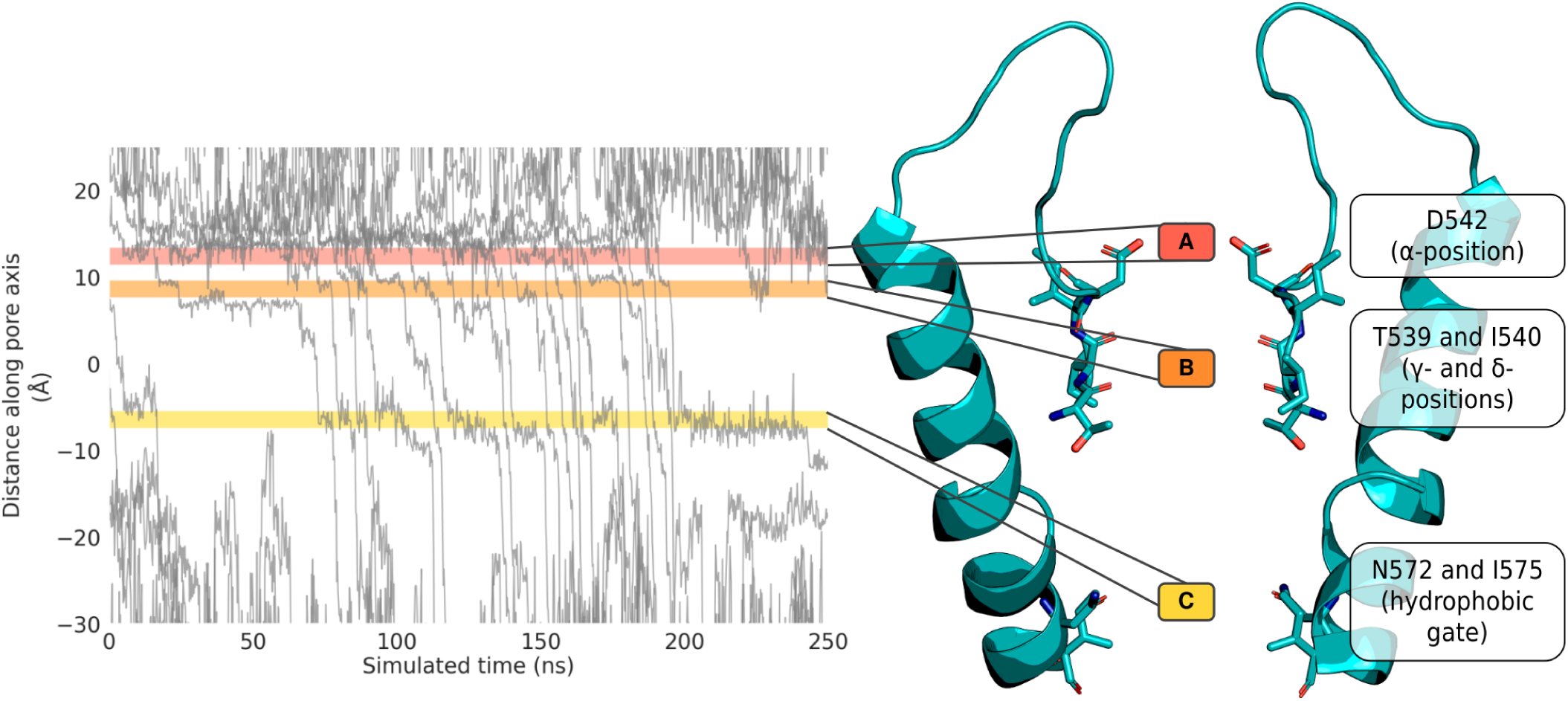
Schematic of cation binding sites identified in the Ca^2+^-selective TRPV5 channel. Permeation traces of the *z*-coordinate of permeating Ca^2+^ cations over time established that cations are bound at three regions within the pore (*left*). The location of the residues constituting these three binding sites in Ca^2+^-selective TRPV channels is shown on the structure of TRPV5 (*right*).

Of the three binding sites we observed, binding site A is formed by the carboxylate oxygen atoms of the ring of acidic residues at the SF entrance (referred to here as the SF *α*-position); binding site B is formed by the carbonyl oxygen atoms of the bottom two SF residues (SF *γ*- and *δ*-positions); and binding site C is formed jointly by the hydrophobic gate consisting of a ring of isoleucine residues (I575 in TRPV5), and by the amide oxygen atoms of the neighbouring asparagine residues (N572 in TRPV5) near the cytoplasmic exit of the pore (Figure 2). The location of these binding sites coincides with constrictions in the pore profile, as determined using CHAP (47) (Figure S1). The distance between binding sites A and B is ∼ 5 Å, and that between binding sites B and C is ∼ 14 Å. We note that Hughes *et al*. (41) reported a further constriction below the hydrophobic gate (binding site C) formed by W583 in the TRPV5 structure, and an analogous constriction at W583 can be observed in TRPV6 (42). However, our simulations do not suggest that the side chains of W583 constitute a functionally important ion binding site, as shown in Figure S3.

In mono-cationic Ca^2+^ solutions, the three binding sites showed Ca^2+^ occupancy probabilities of 0.69 ± 0.05, 0.67 ± 0.04, and 0.57 ± 0.03 in TRPV5, and 0.43 ± 0.04, 0.54 ± 0.05, and 0.29 ± 0.02 in TRPV6 (from A to C, respectively; Figure 3). In mono-cationic Na^+^ solutions, similar occupancies were observed. However, the Na^+^ residence times (*t*_*r*_) at the three binding sites were markedly lower than those observed for Ca^2+^, with ratios of *t*_*r*_(Ca^2+^) : *t*_*r*_(Na^+^) varying between ∼ 35:1 and ∼ 3:1 (Figure 3). These residence times suggest that Ca^2+^ ions have a greater affinity for these binding sites than Na^+^.

**Fig. 3.**
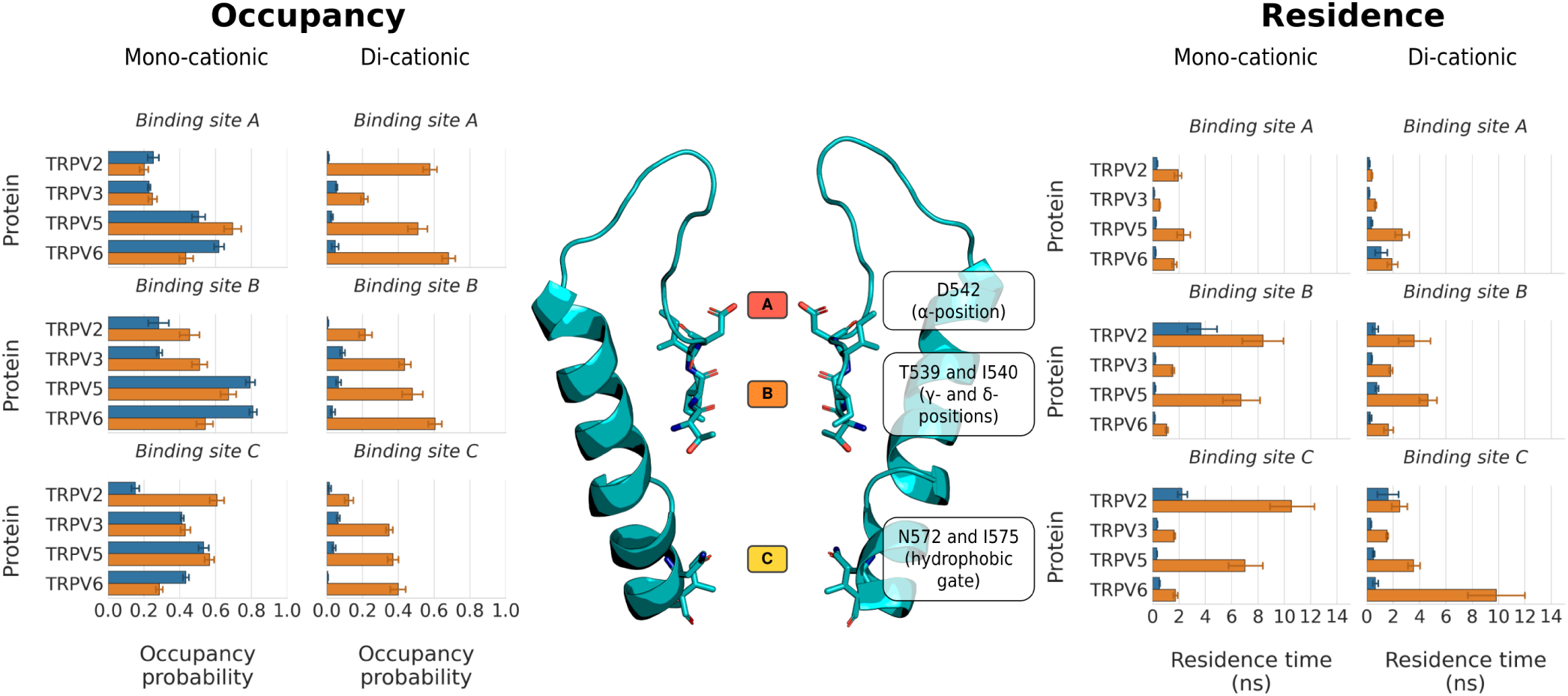
Occupancy probability (*left*) and residence times (*right*) of Ca^2+^ (*orange columns*) and Na^+^ (*blue columns*) cations from simulations of ion permeation in both mono-cationic and di-cationic ion solutions. The mean occupancy probability and SEM were calculated from non-overlapping 50 ns windows from five-fold replicated 250 ns simulations; the mean residence time and SEM from five-fold replicated 250 ns simulations. The location of the residues constituting binding sites A, B, and C in Ca^2+^-selective TRPV channels is shown on the structure of TRPV5 (*centre*).

This observation was further substantiated when the occupancy of binding sites in di-cationic solutions was analysed, in which Ca^2+^ and Na^+^ cations are competing for the binding sites. Binding sites A, B and C in the Ca^2+^-selective channels all showed high occupancies with Ca^2+^ in the mixed cationic solutions (Figure 3). Across all the replicate simulations, we recorded average Ca^2+^ occupancies of 0.51 ± 0.05, 0.48 ± 0.06, and 0.37 ± 0.03 for binding sites A, B and C in TRPV5, respectively, and of 0.68 ± 0.04, 0.61 ± 0.04, and 0.40 ± 0.04 for binding sites A, B and C in TRPV6 (Figure 3). By contrast, the Na^+^ occupancy of each of the three binding sites under these conditions was found to be below 0.07, both in TRPV5 and TRPV6; that is, the ratio between Ca^2+^ and Na^+^ occupancy varies between ∼ 85:1 and ∼ 7:1 (Figure 3). These values indicate a free energy difference of between 11.5 kJ mol^-1^ and 5.0 kJ mol^-1^ for the preferential binding of Ca^2+^ over Na^+^.

### A highly co-operative knock-on mechanism between three cation binding sites underpins selective Ca^2+^ permeation in TRPV channels

The observed increased affinity for Ca^2+^ cations at the pore binding sites compared to Na^+^ means that, in a mixed solution, Ca^2+^ will preferentially occupy these binding sites; however, this also implies that Ca^2+^ ions face a greater energy penalty to dissociate from the binding sites. In mono-cationic solutions, this would result in a greatly reduced Ca^2+^ conductance with respect to Na^+^. For instance, based upon our observed residence times in mono-cationic solutions, we would expect an approximately 12-fold reduced Ca^2+^ unbinding rate compared to Na^+^ for binding site A in TRPV5. However, a much reduced Ca^2+^ conductance is neither observed in our simulations nor in the experimental literature. Due to the divalent charge of Ca^2+^ increasing the affinity to cation binding sites, this dichotomy had previously been suggested to exist, and it was hypothesised that this paradox could be resolved by assuming co-operativity between successive unbinding events such as in a knock-on mechanism (7).

In the classic knock-on mechanism, which for example underpins K^+^ channel function, ions transition into and out of multiple ion binding sites in a highly correlated fashion (48, 49). For example, early experiments by Hodgkin and Keynes and later flux-ratio measurements established that 3–3.4 K^+^ ions moved in lockstep with each other during permeation through K^+^ channels (50, 51).

For each permeating ion in a simulation of TRPV5, Figure 4 shows the association and dissociation of Ca^2+^ and Na^+^ ions at binding sites A, B and C from top to bottom as colour code (bound to A, red; bound to B, orange; bound to C, yellow; transiting within the pore but not bound to a binding site, blue; located in extracellular solvent, dark grey; located in intracellular solvent, light grey). As can be seen for Ca^2+^ in TRPV5 for example (Figure 4 *left*), the plot demonstrates that (i) permeating Ca^2+^ ions spend the vast majority of their time within the pore at the three binding sites (reflected in the scarcity of blue boxes vs. red, orange and yellow), (ii) dual and triple occupancy of the three sites, A, B, and C, with Ca^2+^ is frequently observed (horizontal slices across plot: triple occupancy is observed in 27.2% of the simulation frames, dual occupancy in 49.7%), and (iii) transitions between states show a high degree of correlation, i.e. the ions frequently move in concert into and out of their respective binding sites (horizontal slices; binding state transitions). By contrast, during Na^+^ permeation (Figure 4 *centre*), the ions are predominantly transiting across the pore without occupying particular binding sites for extended time spans (blue, on average 53% of the traversal time for each ion).

**Fig. 4.**
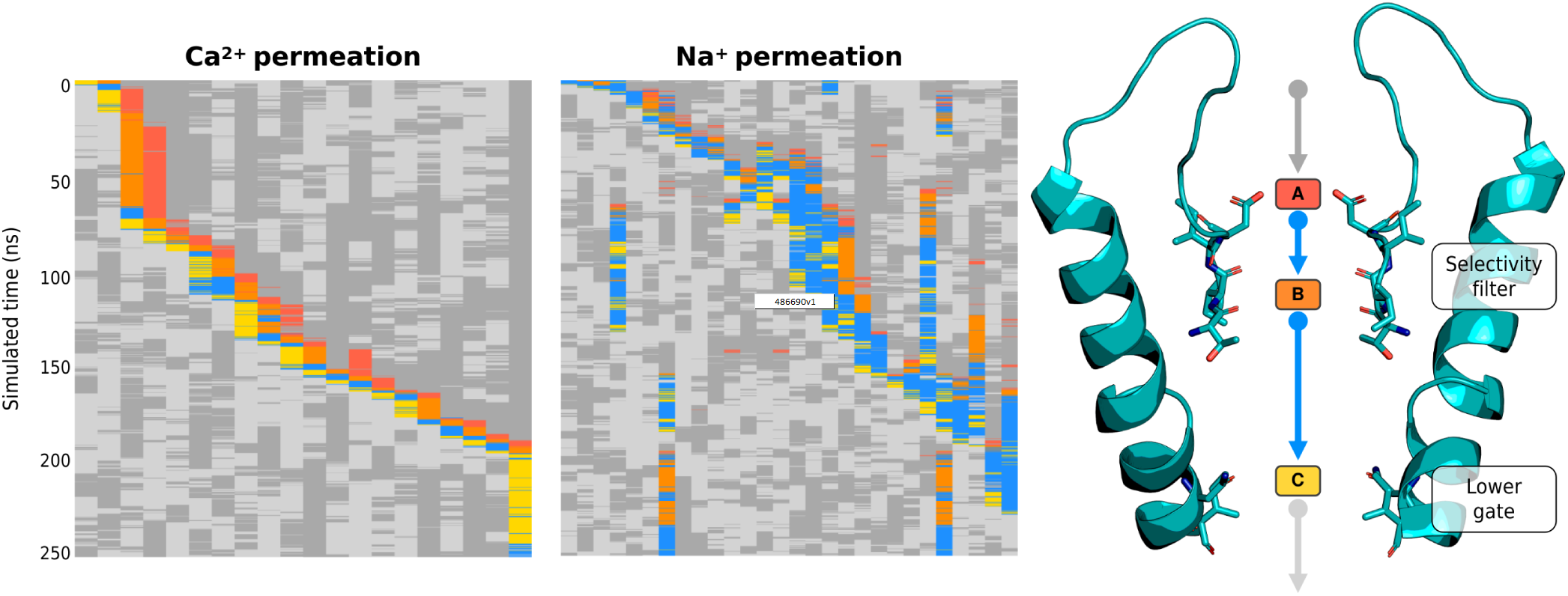
Permeation state plots of permeating Ca^2+^ (*left*) and Na^+^ (*centre*) cations through the Ca^2+^-selective TRPV5. Permeation state plots shows the state of each permeating cation (columns) at a given time point (rows) by assigning a state to the ions to indicate whether the cation is bound to a binding site, transitioning between binding sites, or in the bulk solution: bound to A, red; bound to B, orange; bound to C, yellow; transiting within the pore but not bound to a binding site, blue; located in extracellular solvent, dark grey; located in intracellular solvent, light grey..Comparison of permeation state plots for Ca^2+^ and Na^+^ cations show that Ca^2+^ permeation proceeds in a well-ordered manner with three Ca^2+^ cations within the pore and knocking adjacent cations to the next binding site. Na^+^ permeation on the other hand is far less ordered, with regularly more than three Na^+^ cations within the pore at a given time. Each plot (*left* and *centre*) shows exemplars from a single 250 ns simulation of TRPV5 performed in mono-cationic 150 mM CaCl_2_ or 150 mM NaCl, respectively. The structure of TRPV5 shows the colours used in the permeation state plots and the location of the residues constituting the three binding sites (*right*).

In order to go beyond just visual inspection of the trajectories and to assess the co-operativity of ion permeation in a quantitative way, we developed a new approach based on mutual information, taking into account the “state” of each ion binding site. To achieve this, we assigned a specific binding state (unoccupied, or occupied with a specific ion) to binding sites A, B, and C and used our recently developed approach, state-specific information (SSI, (52)) on pairs of adjacent, permeating cations, to quantify the degree of coupling between ion binding transitions at each of these sites (see *Methods*). This analysis yields a coefficient quantifying the co-operativity between ion binding and unbinding at neighbouring or more distant binding sites, where a greater coefficient signifies a higher degree of coupling; which suggests that when an ion transitions from one site it is more likely there is a transition at the other. To correct for the non-zero mutual information that samples of two completely independent variables can display due to finite-size effects, we followed the approach of McClendon *et al*. (53) and Pethel *et al*. (54) to yield *excess* mutual information, or excess SSI (*exSSI*). We also determined a theoretical upper limit for the maximum mutual information that can be shared between two binding sites by using the minimum state entropy amongst the two sites. Note that this quantity represents an absolute upper limit; reaching it would require both binding sites to exhibit idealised simultaneous states and state transitions throughout the entire simulated time.

The SSI analysis showed that in the Ca^2+^-selective TRPV channels, TRPV5 and TRPV6, a high level of information above noise is shared between the transition of ions into and out of binding sites A and B, respectively, both for the permeation of Ca^2+^ and Na^+^ (*exSSI* between 0.8–1.6 bits, Table S5 and Figure 5). This suggests that the ion binding and unbinding processes at each of these binding sites are coupled with one another, constituting a knock-on mechanism at relatively short range.

**Fig. 5.**
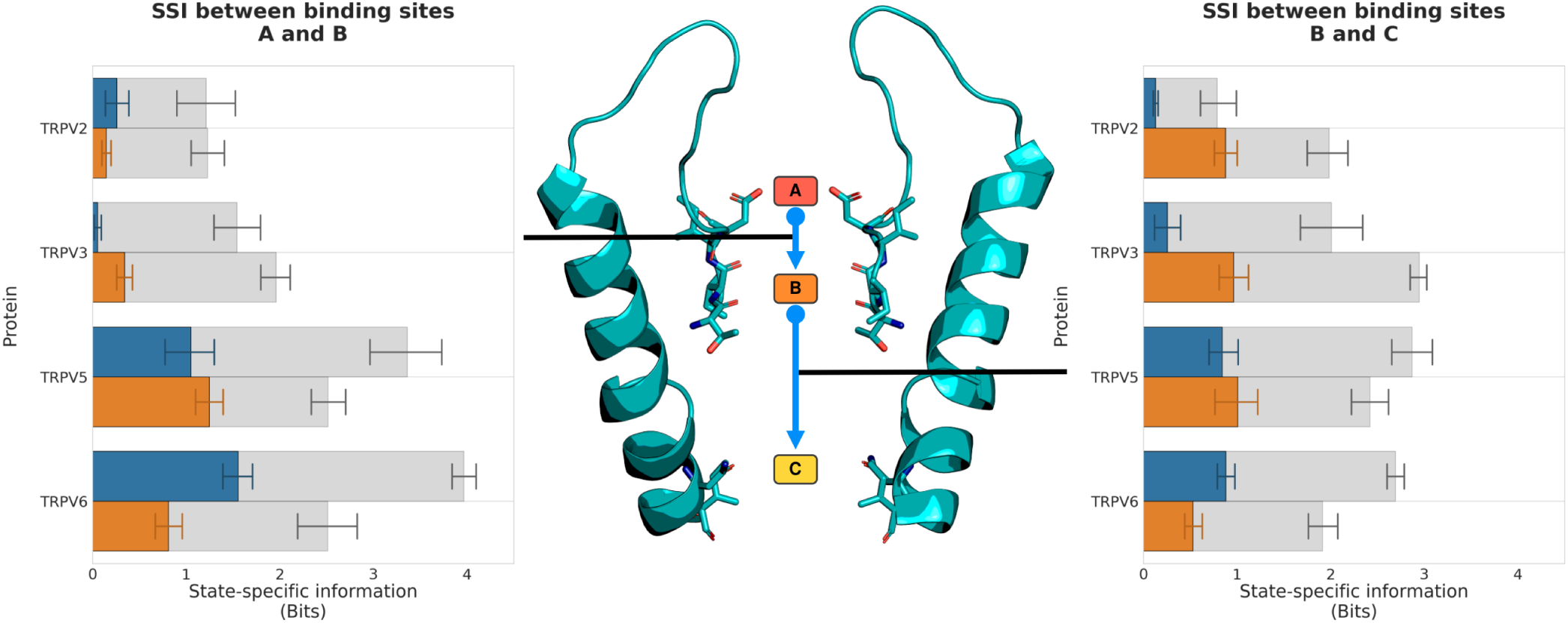
Excess state-specific information (*exSSI*) between ion binding sites quantifies the degree of co-operativity in the knock-on mechanism of cation permeation in TRPV channels. The mean *exSSI* and SEM between transitions from binding sites A and B (*left*) and binding sites B and C (*right*) are shown for Ca^2+^ (*orange*) and Na^+^ (*blue*) cations in mono-cationic solutions. For each *exSSI*, the mean maximum *exSSImax* and standard error is also shown (*grey*), and the *exSSInorm* is reported in Table S5.

Similarly, the transitions of ions into and out of binding sites B and C show a large degree of correlation for both Ca^2+^ and Na^+^ (Figure 5). In the case of binding sites B and C however, this requires a remote knock-on mechanism to be in operation, since these sites are ∼ 14 Å apart. The concept of a remote knock-on event was first proposed by Tindjon *et al*. based upon Brownian dynamics simulations (55), and observed by Zhang *et al*. in atomistic MD simulations of Ca^2+^ permeation in the RyR1 channel (39). Our SSI analysis suggests that the degree of co-operativity in the remote knock-on mechanism between binding sites B and C (Figure 5 *left*) is slightly smaller than the co-operativity in the direct knock-on mechanism between binding sites A and B (Figure 5 *right*).

### Cation permeation in non-selective TRPV channels shows a lower degree of co-operativity

To determine if the remaining, non-selective TRPV channels showed a different permeation mechanism, we next performed simulations of the open-state TRPV2 (56) and TRPV3 (57) channels using the same simulation approach as described for the Ca^2+^-selective TRPV channels. These simulations of non-selective TRPV channels also showed continuous ion permeation, with cation conductances, again, in good agreement with published conductance values measured *in vitro* (Table 2). Overall, we recorded 706 complete inward channel crossings for Ca^2+^ and 1,176 for Na^+^ from simulations of the non-selective TRPV channels.

**Table 2.** Calculated conductances from MD simulations of ion permeation in non-selective TRPV channels. Mean inward conductances and standard error of mean (SEM) were calculated from overlapping 50 ns windows from five-fold replicated 250 ns simulations. Mono-cationic solutions at 150 mM concentration; di-cationic mixture of Ca^2+^ and Na^+^ at a concentration of 75 mM each.

|  | Conductance (pS) |  |  |  |
| --- | --- | --- | --- | --- |
| | $\text{Ca}^{2+}$ | $\text{Na}^+$ | $\text{Ca}^{2+}$ & $\text{Na}^+$ | Literature |
| TRPV2 | 40 ± 4 | 18 ± 3 | 43 ± 7 | 28 ($\text{Na}^+$) (58) |
| TRPV3 | 196 ± 10 | 299 ± 10 | 222 ± 13 | 201 ($\text{Na}^+$) (59) |

The occupancy of the three binding sites in the TRPV2 and TRPV3 systems showed no clear difference to the Ca^2+^-selective channels; however the occupancy of binding site A was slightly reduced for both cations in the mono-cationic solutions (Figure 3). All binding sites exhibited a preference for binding Ca^2+^ in the di-cationic solutions.

We were therefore curious if the three-site knock-on mechanism described previously for Ca^2+^-selective TRPV channels is also at play in the non-selective TRPV channels. Our SSI analysis confirmed that the co-operativity between binding sites B and C in the non-selective TRPV channels was comparable to those calculated for the Ca^2+^-selective TRPV channels (Table S5 and Figure 5). However, the correlation between ion binding transitions at binding sites A and B was substantially reduced in both of the non-selective TRPV systems (Figure 5). The nearly complete absence of co-operativity from binding sites A and B demonstrates that a knock-on mechanism is not occurring between these two sites in the non-selective TRPV channels.

The slightly reduced occupancy at binding site A, coupled with the loss of co-operativity between ion binding transitions at binding sites A and B, suggests that in non-selective TRPV channels binding site A is not able to optimally coordinate incoming cations in a way that it constitutes part of the knock-on mechanism of cation permeation. Instead, our findings suggest that cation permeation in the non-selective TRPV channels occurs via a two-site knock-on mechanism between binding sites B and C.

Based on our SSI approach to quantify mutual information in binding and unbinding events at different ion binding sites, and using the concept of *total correlation* to evaluate the overall co-operativity in a system across all coupled events, we next calculated the total correlation of ion permeation for all the TRP channels investigated. The reduction in the number of correlated knock-on sites within the non-selective TRP channels can be distinctly seen when comparing the total correlation for each of the non-selective and Ca^2+^-selective TRPV channels (Figure S4).

The ion binding sites in non-selective TRPV channels preferentially bind Ca^2+^ over Na^+^, as in Ca^2+^-selective TRPV channels, which explains why the non-selective TRPV channels, in fact show slight Ca^2+^-selectivity (*P*_*Ca*_*/P*_*Na*_ ∼ 10:1 (28)). However, our data suggests that the reduced level of coordination at binding site A, and especially the effect on the three-site co-operativity it imparts together with sites B and C, reduces the Ca^2+^-selectivity from *P*_*Ca*_*/P*_*Na*_ ∼ 100:1 seen in TRPV5 and TRPV6, and in this way ultimately determines the difference between Ca^2+^-selective and non-selective permeation.

We note that the *P*_*Ca*_*/P*_*Na*_ values obtained from our simulations overall show lower Ca^2+^ selectivity than the reported literature values (Table S6). We surmised that this might be due to the higher voltages used in our simulations to enhance the sampling rate. Supplementary simulations performed at a lower voltage demonstrated that, indeed, the selectivity for Ca^2+^ increases with lower voltages across the membrane (Figure S5). Below a voltage threshold of ∼ 205 mV, however, the sampling of permeation events in the simulations became very poor, such that we were not able to reliably probe the precise voltage range used in the experiments.

### Structural features distinguishing Ca^2+^-selective from non-selective permeation

To understand why cation coordination at binding site A is weakened in non-selective TRPV channels, decoupling its co-operativity, we investigated the area formed between the four subunits of the TRPV channel for each residue in the selectivity filter. The cross-sectional area formed by the carboxylate oxygen atoms at the SF *α*-position is larger in the non-selective TRPV channels than in the Ca^2+^-selective TRPV channels (Figure 6). Interestingly, despite the increased average area formed by the carboxylate oxygen atoms at the *α*-position residue in the selectivity filter, the average area formed by the carbonyl oxygen atoms at both the *γ*- and *δ*-positions are smaller in non-selective TRPV channels than in Ca^2+^-selective TRPV channels (Figure 6). These differences in selectivity architecture were further confirmed using the pore profile calculated using CHAP (47) (Figure S1). Our simulations showed no differences in the flexibility of selectivity filters between Ca^2+^-selective and non-selective TRPV channels as determined by RMSF calculations of the backbone atoms (Table S7); we did however observe a small difference in the backbone dihedral angle distribution of the *β*-position residue, which we predict to be due to differences in stabilising hydrophobic interactions of the residue side-chain (Figure S6).

**Fig. 6.**
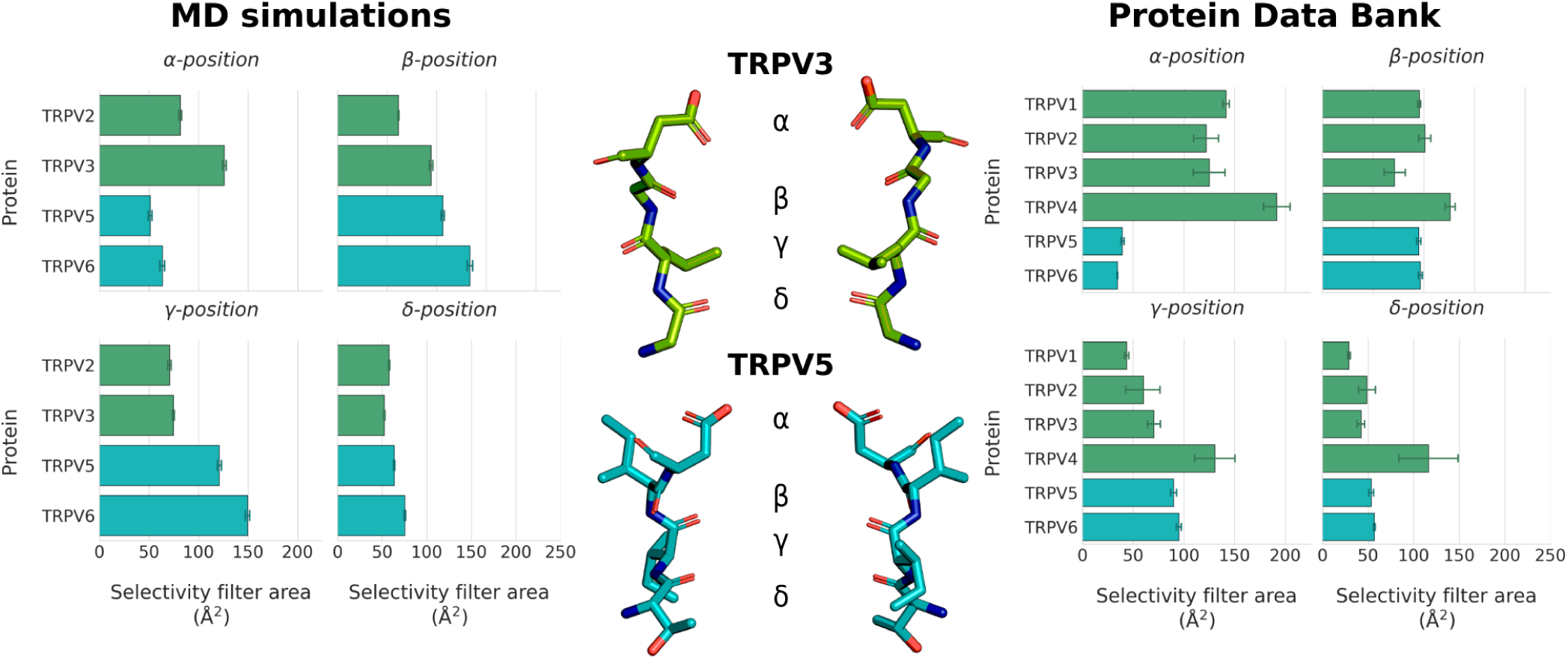
Architecture of the four-residue selectivity filter of the investigated TRPV channels. The area formed between the carboxylate oxygen atoms (*α*-position) and backbone carbonyl atoms (*β*-, *γ*-, and *δ*-positions) was calculated for all Ca^2+^-selective and non-selective TRPV systems simulated as part of this study (*left*), and for all TRPV structures available in the Protein Data Bank at the time of writing (*right*). The SF structures of the Ca^2+^-selective TRPV5 and non-selective TRPV3 are shown for reference (*centre*). The mean area between selectivity filter residues and SEM were calculated from non-overlapping 50 ns windows from five-fold replicated 250 ns simulations in 150 mM CaCl_2_. The mean area between selectivity filter residues and SEM from PDB structures were calculated from all available homotetrameric TRPV structures at the time of writing.

To expand the geometric analysis to all available TRPV channel structures, we also calculated the average area formed by selectivity filter residues from all available TRPV structures deposited in the Protein Data Bank (PDB) (Figure 6). Analysis of these static structures showed the same two trends observed within our MD simulations: (1) Ca^2+^-selective TRPV channels clearly have a smaller average area formed by the carboxylate oxygen atoms of the SF *α*-position; and (2), non-selective TRPV channels have a slightly narrower constriction at the SF *γ*- and *δ*-position residues. Our MD simulations suggested that the wider opening at the *α*-position leads to a weaker cation binding interaction at binding site A and cation coordination, which, in turn, decouples binding site A from the co-operative knock-on mechanism that underpins permeation in the Ca^2+^-selective TRPV channels.

### Ca^2+^-selective permeation does not depend on the solvation states of permeation cations

A previously reported mechanism of how cation selectivity can be achieved is by desolvation of permeating cations. Differences in desolvation energies between cationic species provide a thermodynamic penalty that can be more favourable for the permeation of one cationic species over another. Such a mechanism has been reported to underpin K^+^-selectivity over Na^+^ in K^+^ channels (29, 30), for example.

To investigate whether desolvation was a factor in Ca^2+^-selectivity in TRPV channels, we determined the number of oxygen atoms within a 3 Å radius of the cations, representing their first solvation shell (Figure 7). In the bulk solution, both Ca^2+^ and Na^+^ cations showed their expected water co-ordination number of 7 and 5.6, respectively. As the cations entered the pore, we saw a small degree of partial dehydration of permeating cations at the SF (Figure 7. In particular, the carboxylate oxygen atoms of the acidic residue at the entrance of the SF co-ordinated an incoming cation, with these displacing up to ∼ 2 co-ordinated water molecules from the first solvation shell of the cation. We also observe some desolvation around, or below, binding site C. However, no major differences in the solvation shell of permeating Ca^2+^ or Na^+^ cations, or indeed between the Ca^2+^-selective and non-selective TRPV channels were observed. Therefore, this finding suggests that differences in cation desolvation are not a major factor underpinning the differences in Ca^2+^-selectivity between Ca^2+^-selective and non-selective TRPV channels.

**Fig. 7.**
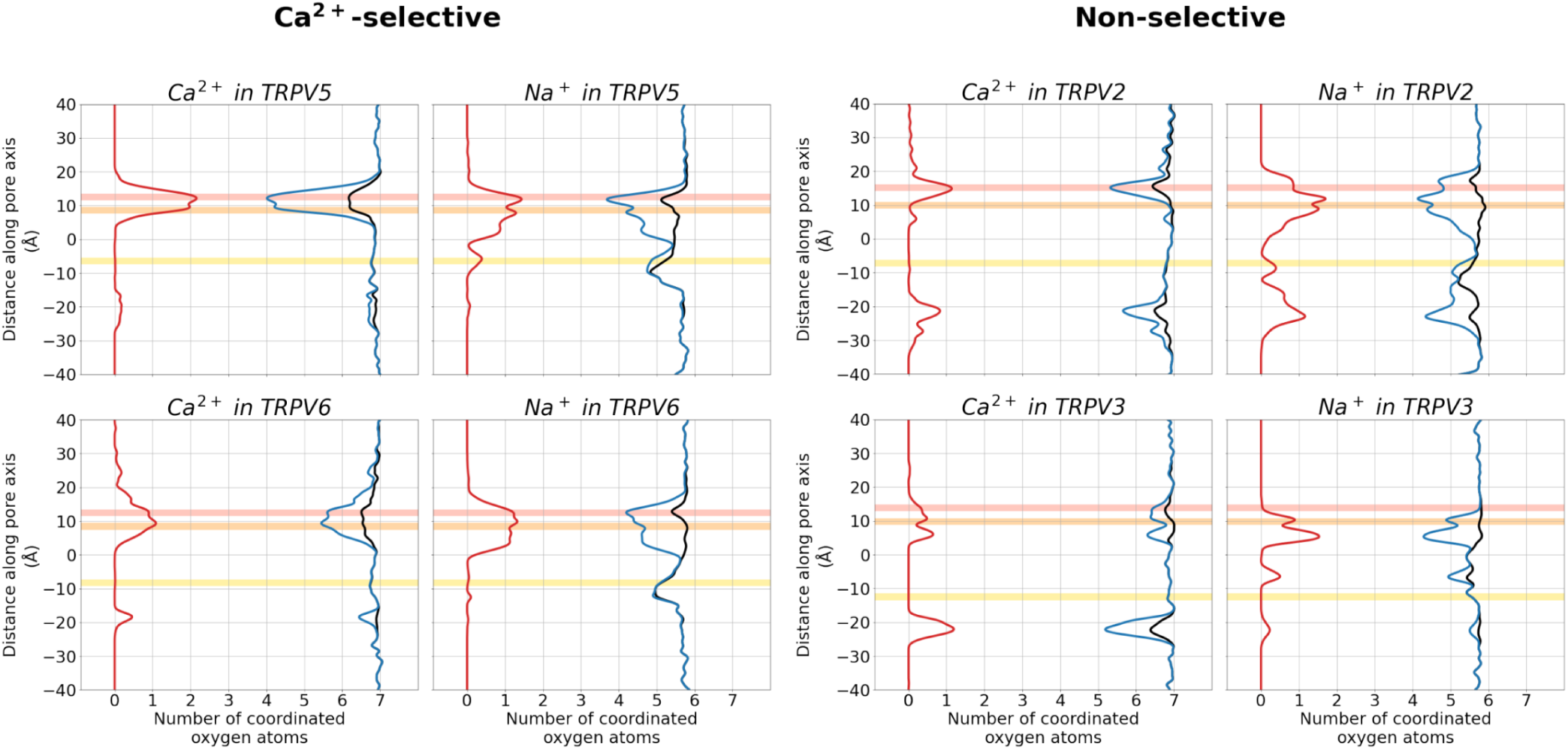
Solvation state of permeating cations as they permeate through TRPV channels. The mean number of oxygen atoms of water molecules (*blue*), the number of oxygen atoms of protein residues (*red*), and total number of any oxygen atoms (*black*) within 3 Å of each permeating cation are plotted. The curves were smoothed using a Gaussian filter with a sigma value of 3.

## Discussion

Our simulations showed three main cation binding sites in the permeation pathway of TRPV channels, which we term binding sites A, B and C. Binding sites A and B are formed by the carboxylate oxygen atoms of the *α*-position residue of the SF and by the carbonyl oxygen atoms of the *γ*- and *δ*-position residues of the SF, respectively. Binding site C is located just above the hydrophobic lower gate of the pore, and is formed by the isoleucine residues of the lower gate and amide oxygen atoms of the neighbouring asparagine residues.

The identification of these cation binding sites in our MD simulations is in agreement with previously published TRPV structures and other MD simulations. In their crystal structure of the *Rattus norvegicus* TRPV6, Saotome *et al*. identified three regions of electron density within the channel pore which they interpreted as cation-binding sites (45). It should be noted that there are some minor differences in the residues forming binding site C to the binding site reported here, likely due to the structure of TRPV6 of Saotome *et al*. being in the closed-state (45), rather than the open-state structure of McGoldrick *et al*. used in the present study (42). During the transition from a closed-state to the open-state, TRP channels undergo a rotation about the pore-forming S6 helix which changes the pore-facing residues (42).

Moreover, numerous structures of TRPV channels deposited within the PDB are resolved with cations bound at one of the cation binding sites identified in our MD simulations. These include structures from several orthologues of the TRPV channels simulated in this study, as well as of the TRPV1 and TRPV4 channels that were not simulated in this study. For example, structures of the closed-state TRPV1 channel of *Rattus norvegicus* solved at 4 °C (60) and the open-state TRPV4 channel of *Homo sapiens* in complex with 4*α*-PDD (61), both model cations bound at binding site B. This observation further validates the existence of the identified cation binding sites, as well as their conservation among TRPV channels and perhaps the wider TRP superfamily.

In addition to structural data, Sakipov *et al*. performed equilibrium MD simulations of Ca^2+^ movement in the closed-state structure of *Rattus norvegicus* TRPV6 and identified cation binding sites in the SF (46). The binding sites from this work are also generally in agreement with our simulations results and the aforementioned structural data. However, Sakipov *et al*. reported that their simulation data identified two Ca^2+^ cations residing at binding site A, with adjacent chains each occupying an ion. In accordance with previous X-ray crystallographic data, two ions associated to binding site A are not seen in a major population of our simulation ensembles, which we attribute to the use of an optimised multi-site model for Ca^2+^ ions in the present study (39) and increased sampling for improved statistical analysis.

In mono-cationic solutions, we observed a high probability of at least two of the three binding sites (A, B and C) being simultaneously occupied with either Na^+^ or Ca^2+^, respectively, with mostly insignificant differences between the occupancy values for Na^+^ and Ca^2+^ at each individual binding site. However, the residence times were markedly reduced at all sites for Na^+^ ions, and overall a tendency towards weaker binding for either ion at binding site A at the extracellular SF entrance was observed.

In mixed, di-cationic solutions of Na^+^ and Ca^2+^, by contrast, Ca^2+^ ions strongly out-competed Na^+^ ions for association at all pore binding sites, both in the Ca^2+^-selective and non-selective TRPV channels. Therefore, according to our data, all the binding sites display a much greater affinity for Ca^2+^. This gave rise to the question of how permeation efficiency for conducting Ca^2+^ ions is achieved in these channels and how this relates to their varying degrees of selectivity, since higher affinity binding is usually expected to lead to slower permeation rates.

By ensuring co-operativity between binding/unbinding events at multiple binding sites, permeation rates can be enhanced (7). We hypothesised that the level of co-operativity between sites A, B and C in the pore could underpin the difference between highly and less Ca^2+^-selective TRPV channels. We therefore developed a novel method to quantify co-operativity during ion permeation in pores with multiple ion binding sites based on mutual information and total correlation measures using the SSI approach (52). We anticipate that this method will be similarly useful for the study of permeation mechanisms and the basis of selectivity in other channels. Our analysis showed that there is a substantial degree of co-operativity for Ca^2+^ permeation between binding sites B and C across all TRPV channels, whereas a clear distinction exists between the co-operativity between binding sites A and B in Ca^2+^-selective and non-selective TRPV channels. In the non-selective TRPV channels, binding site A is decoupled from binding site B. By contrast, binding sites A and B are even more strongly coupled than binding sites B and C in the case of the Ca^2+^-selective channels. We suggest that this marked difference in co-operativity mechanistically explains the different levels of Ca^2+^ selectivity in TRPV channels.

## Conclusions

We have characterised the cation permeation mechanisms in four members of the TRPV channel family. We identified three cation binding sites within the pore, each of which displayed greater affinity for Ca^2+^ binding over Na^+^. A novel application of mutual information between ion binding and unbinding at consecutive binding sites (SSI) enabled us to quantify the degree of knock-on taking place in ion permeation. This showed that the level of Ca^2+^ selectivity in TRPV channels is determined by the co-operativity, or coupling, between the transitions at three pore cation binding sites. The Ca^2+^-selective TRPV channels display a highly correlated three-site knock-on, whereas the cation binding site at the extracellular entrance is decoupled from the mechanism in the non-selective TRPV channels, which reduces the overall preference for Ca^2+^ permeation.

## Methods

### TRPV system construction

Truncated TRPV simulation systems consisting of the membrane-domain of the channels were constructed as described in Table 3. The systems were built using the CHARMM-GUI server (62). The charged *N*- and *C*-terminal residues were neutralised by capping with acetylated (ACE) and *N*-methylamidated (CT3) groups, respectively. All missing non-terminal residues were modelled, and mutations input as required (63). In the case of the TRPV5 system, the parameters for PI(4,5)P_2_ were generated using the CHARMM General Force Field (CGenFF) (64) through the ligand reader and modeller in CHARMM-GUI (65).

**Table 3.** Summary of protein constructs used in this study.

| Protein | PDB ID | Orthologue | Residue range | Upper-gate residue | Lower-gate residue | Reference |
| --- | --- | --- | --- | --- | --- | --- |
| TRPV2 | 6BO4 | *Rattus norvegicus* | 327 to 691 | E609 | I642 | (56) |
| TRPV3 | 6PVP | *Mus musculus* | 375 to 745 | D641 | I674 | (57) |
| TRPV5 | 6DMU | *Oryctolagus cuniculus* | 262 to 639<br>(and PI(4,5)P<sub>2</sub>) | D542 | I575 | (41) |
| TRPV6 | 6BO8 | *Homo sapiens* | 262 to 638 | D542 | I575 | (42) |

The structures were aligned in the membrane using the PPM server (66), and inserted into a 1-palmitoyl-2-oleoyl-sn-glycerol-3-phosphocholine (POPC) bilayer of 150 × 150 Å size using the CHARMM-GUI membrane builder (67, 68), and then solvated. Ions were added using GROMACS 2020.2 (69, 70) to neutralise any system charges and add ions to a concentration of either 150 mM NaCl, 150 mM CaCl_2_, or a mixture of 75 mM NaCl and 75 mM CaCl_2_. In the case of simulations containing Ca^2+^, the standard CHARMM36m Ca^2+^ molecules were then replaced with the multi-site Ca^2+^ of Zhang *et al*. (39). Hydrogen mass re-partitioning (HMR) of the system was used to allow the use of 4-fs time steps in simulations of NaCl solutions. The multi-site Ca^2+^ model used for simulations of CaCl_2_ however is incompatible with a 4-fs time step, and therefore any simulations including Ca^2+^ cations were performed with HMR but at a time step of 2-fs. The protein was restrained in the open-state by applying harmonic restraints on the *α*-carbon atoms of the lower gate residues (see Table 3).

### Molecular dynamics simulations details

All simulations were performed using GROMACS 2020.2 (69, 70) and the CHARMM36m force field for the proteins, lipids, and ions (71). The TIP3P water model was used to model solvent molecules (72). The system was minimised and equilibrated using the suggested equilibration inputs from CHARMM-GUI (73). In brief, the system was equilibrated using the NPT ensemble for a total time of 1.85 ns with the force constraints on the system components being gradually released over six equilibration steps. The systems were then further equilibrated by performing a 15 ns simulation with no electric field applied. To prevent closing of the lower-gate of the pore, harmonic restraints were applied to maintain the distance between the *α*-carbon atoms of the lower gate residues of each respective chain (Table 3). To drive ion permeation, an external electric field was applied by using the method of Aksimentiev *et al*. (74) to production simulations with an E_0_ of -0.03 V nm^-1^; this resulted in a transmembrane voltage of ∼410 mV with negative polarity in the intracellular region. The temperature was maintained at 310 K using the Nose-Hoover thermostat (75) and the pressure was maintained semi-isotropically at 1 bar using the Parrinello-Rahman barostat (76). Periodic boundary conditions were used throughout the simulations. Long-range electrostatic interactions were modelled using the particle-mesh Ewald method (77) with a cut-off of 12 Å. The LINCS algorithm (78) was used to constrain bonds with hydrogen atoms. All individual simulations were 250 ns long and repeated five times for each system (Tables S1 and S2), with the exception of the additional control simulations described in Table S3.

### Simulation analysis

Analysis of MD trajectory data was performed using in-house written Python scripts, utilising GROMACS modules (69, 70), the SciPy library of tools (79–82), and MDAnalysis (83, 84). Analysis of the pore architecture was performed using CHAP (47). All plots were generated in Python using Matplotlib (85) and Seaborn (86). All MD inputs and analysis scripts used for this study are deposited in a public GitHub repository, available at: https://github.com/cmives/Ca_selectivity_mechanism_of_TRPV_channels.

### Calculating conductance and selectivity from in silico electrophysiology experiments

The conductance of the channels (*C*_*ion*_) was calculated according to Equation 1, where *N*_*p*_ is the number of permeation events, *Q*_*ion*_ is the charge of the permeating ion in Coulomb, *t*_*traj*_ is the length of the trajectory, and *V*_*tm*_ is the transmembrane voltage. The mean conductance and standard error were calculated from overlapping 50 ns windows of the trajectory.

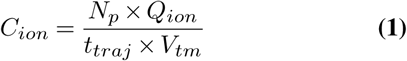

The selectivity (*P*_*Ca*_*/P*_*Na*_) was calculated as the ratio between the total sum of Ca^2+^ permeation events and the total sum of Na^2+^ permeation events, from five-fold replicated 250 ns simulations in di-cationic solutions with 75 mM CaCl_2_ and 75 mM NaCl.

### Identification of cation binding sites from MD simulations of TRPV channels

Cation binding sites were identified by plotting timeseries of each permeating ion with respect to their position along the pore axis. To further validate this, the *Featurizer* function of PENSA (44) was used to identify the 12 most occupied ion binding sites, as determined by 3D density maxima within 7 Å of the protein. This analysis was performed on a trajectory of concatenated five-fold replicated 250 ns simulations with a 200 ps time-step from mono-cationic simulations.

### Calculating ion occupancy probabilities and residence times in the identified cation binding sites

The ion occupancy (*Oion*) of the identified cation binding sites was calculated by dividing the number of frames (*N*_*occupied*_), in which an ion’s centre of geometry is within 3.5 Å of the centre of geometry of the ion co-ordinating binding site atoms by the number of frames in the time window (*N*_*frames*_). The mean ion occupancies and standard error were calculated from non-overlapping 50 ns windows of the five-fold replicated 250 ns simulation trajectories with a 20 ps time-step.

The ion residence times (*t*_*r*_) were calculated by averaging the amount of time an individual ion was located within 3.5 Å of the centre of geometry of the ion co-ordinating binding site atoms. The mean *t*_*r*_ and standard error were calculated from five-fold replicated 250 ns simulation trajectories with a 20 ps time-step.

### Characterising permeation co-operativity through mutual information using SSI from PENSA

To characterise the level of co-operativity in the knock-on permeation mechanisms in TRPV channels, we used PENSA to calculate the state-specific information (SSI) shared between discrete state transitions in the occupancy distributions of each binding site (44, 52).

A timeseries distribution with a timestep of 20 ps for each binding site was obtained, whereby for each frame, if an ion occupied the binding site then this ion’s atom ID number was recorded, whereas if the binding site was unoccupied, an ID of -1 was recorded. The ID numbers were discrete, and changes between ID numbers in each binding site therefore represent discrete state transitions. By quantifying the information shared between changes to the ID numbers in each site, we were able to determine whether ion transitions at one site were coupled to transitions at another during a 20 ps time interval. From this we could conclude whether cations are “knocking” each other. Furthermore, we employed a custom filter to separate binding site data into specific ion types, which allowed us to calculate whether binding site co-operativity depended on the types of ions transitioning through the sites in simulations with a di-cationic solution.

Similar to McClendon *et al*., we observed that finite sampling resulted in independent distributions sharing mutual information (53, 54). To overcome this, we calculated a statistical threshold for each simulation via randomly permuted copies of the original data. Random permutations of the original data maintained marginal probabilities for each simulation while at the same time quantifying the effect of finite sampling on the measurement of state-specific information. Since the upper bound of mutual information between two variables is equal to the lowest entropy of those variables, we used the binding site corresponding to the lowest entropy for obtaining the threshold. This ensured that the portion of SSI which could be attributed to random noise between any two binding sites was always less than or equal to the SSI. State-specific information was then calculated between two independently permuted versions of the occupancy distribution for the minimum entropy binding site. This measurement was repeated 1,000 times in order to resolve a Gaussian distribution from which we obtained the 99% confidence threshold. We subtracted this threshold from the measured values to resolve excess mutual information, or excess SSI (*exSSI*), shared in discrete state transitions. As it is not possible to have negative information transfer, negative *exSSI* values were masked to a value of 0. We also derived a maximum SSI value representing a theoretical upper limit for the information that can be shared between two binding sites, where *exSSI*_*max*_ is given by subtracting the random threshold from the minimum entropy of the two binding sites in question.

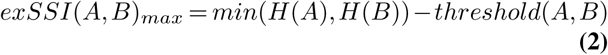

To quantify the interdependency of all three ion binding sites within the TRPV pores, the total correlation (TotCorr) was obtained using Equation 3, where *H*(*A*), *H*(*B*), *H*(*C*) represent the entropy of binding sites *A, B* and *C*, respectively, and *H*(*A, B, C*) the joint entropy of binding sites *A, B* and *C*.

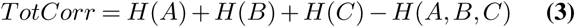

### Characterising the architecture of the selectivity filter of TRPV channels

To determine the area formed between residues in the SF, the area of the quadrilateral between the adjacent chains was calculated on the *X* and *Y* axes. For this, we used the carboxylate oxygen atoms of the adjacent chains for the *α*-position residue, and the carbonyl oxygen atoms for the *β*-, *γ*-, and *δ*-position residues.

To quantify the SF areas from our MD simulations, the mean and standard error were calculated from non-overlapping 50 ns windows of the five-fold replicated 250 ns trajectories with a 200 ps time-step. Furthermore, the SF areas of all TRPV structures deposited in the PDB, available as of 4^th^ February 2022, were determined. A total of 101 structures were analysed, with non-tetrameric structures or structures without all the atoms of interest modelled not included. The mean and standard error of the mean were calculated for all the available structures for any particular TRPV channel.

## ACKNOWLEDGEMENTS

We thank Professor Chen Song of Peking University for providing parameters and discussion on the multi-site Ca^2+^ model. We also thank the University of Dundee I.T. services for maintenance of the School of Life Sciences high-performance computing (HPC) cluster which is utilised in this research. CMI was supported by the Medical Research Council [grant number MR/N013735/1], and NJT was supported by the UKRI Biotechnology and Biological Sciences Research Council (BBSRC) [grant number BB/M010996/1].

## Supplementary information

### Summary of MD simulations within this study

Tables S1, S2, and S3 show details of all MD simulations performed in this study.

**Table S1.** Summary of simulation details of Ca^2+^-selective TRPV channels.

| Protein | TRPV5 |  |  | TRPV6 |  |  |
| --- | --- | --- | --- | --- | --- | --- |
| Structure | 6DMU (41)<br>(262-639 & PI(4,5)P<sub>2</sub>) |  |  | 6BO8 (42)<br>(262-638) |  |  |
| Force field | CHARMM36m (71) |  |  | CHARMM36m (71) |  |  |
| Water | TIP3P (72) |  |  | TIP3P (72) |  |  |
| Ligand | PI(4,5)P<sub>2</sub> (CGenFF (64)) |  |  | - |  |  |
| Ion | 150 mM CaCl<sub>2</sub><br>291 Ca<sup>2+</sup> (Zhang *et al.* (39))<br>574 Cl<sup>-</sup> (CHARMM36m (71)) | 150 mM NaCl<br>295 Na<sup>+</sup> (CHARMM36m (71))<br>287 Cl<sup>-</sup> (CHARMM36m (71)) | 75 mM CaCl<sub>2</sub> + 75mM NaCl<br>143 Ca<sup>2+</sup> (Zhang *et al.* (39))<br>151 Na<sup>+</sup> (CHARMM36m (71))<br>429 Cl<sup>-</sup> (CHARMM36m (71)) | 150 mM CaCl<sub>2</sub><br>279 Ca<sup>2+</sup> (Zhang *et al.* (39))<br>562 Cl<sup>-</sup> (CHARMM36m (71)) | 150 mM NaCl<br>279 Na<sup>+</sup> (CHARMM36m (71))<br>283 Cl<sup>-</sup> (CHARMM36m (71)) | 75 mM CaCl<sub>2</sub> + 75mM NaCl<br>140 Ca<sup>2+</sup> (Zhang *et al.* (39))<br>140 Na<sup>+</sup> (CHARMM36m (71))<br>424 Cl<sup>-</sup> (CHARMM36m (71)) |
| Independent simulations | 5 | 5 | 5 | 5 | 5 | 5 |
| Total simulation time (μs) | 1.25 | 1.25 | 1.25 | 1.25 | 1.25 | 1.25 |
| Estimated voltage (mV) | -410 | -410 | -410 | -410 | -410 | -410 |
| Permeation events | 85 | 165 | 165 | 189 | 197 | 49 |

**Table S2.** Summary of simulation details of non-selective TRPV channels.

| Protein | TRPV2 |  |  | TRPV3 |  |  |
| --- | --- | --- | --- | --- | --- | --- |
| Structure | 6BO4 (56)<br>(327-691) |  |  | 6PVP (57)<br>(375-745) |  |  |
| Force field | CHARMM36m (71) |  |  | CHARMM36m (71) |  |  |
| Water | TIP3P (72) |  |  | TIP3P (72) |  |  |
| Ligand | - |  |  | - |  |  |
| Ion | 150 mM CaCl<sub>2</sub><br>253 Ca<sup>2+</sup> (Zhang *et al.* (39))<br>526 Cl<sup>-</sup> (CHARMM36m (71)) | 150 mM NaCl<br>253 Na<sup>+</sup> (CHARMM36m (71))<br>273 Cl<sup>-</sup> (CHARMM36m (71)) | 75 mM CaCl<sub>2</sub> + 75mM NaCl<br>127 Ca<sup>2+</sup> (Zhang *et al.* (39))<br>127 Na<sup>+</sup> (CHARMM36m (71))<br>401 Cl<sup>-</sup> (CHARMM36m (71)) | 150 mM CaCl<sub>2</sub><br>291 Ca<sup>2+</sup> (Zhang *et al.* (39))<br>582 Cl<sup>-</sup> (CHARMM36m (71)) | 150 mM NaCl<br>291 Na<sup>+</sup> (CHARMM36m (71))<br>291 Cl<sup>-</sup> (CHARMM36m (71)) | 75 mM CaCl<sub>2</sub> + 75mM NaCl<br>146 Ca<sup>2+</sup> (Zhang *et al.* (39))<br>146 Na<sup>+</sup> (CHARMM36m (71))<br>438 Cl<sup>-</sup> (CHARMM36m (71)) |
| Independent simulations | 5 | 5 | 5 | 5 | 5 | 5 |
| Total simulation time (μs) | 1.25 | 1.25 | 1.25 | 1.25 | 1.25 | 1.25 |
| Estimated voltage (mV) | -410 | -410 | -410 | -410 | -410 | -410 |
| Permeation events | 60 | 59 | 72 | 941 | 309 | 441 |

**Table S3.**
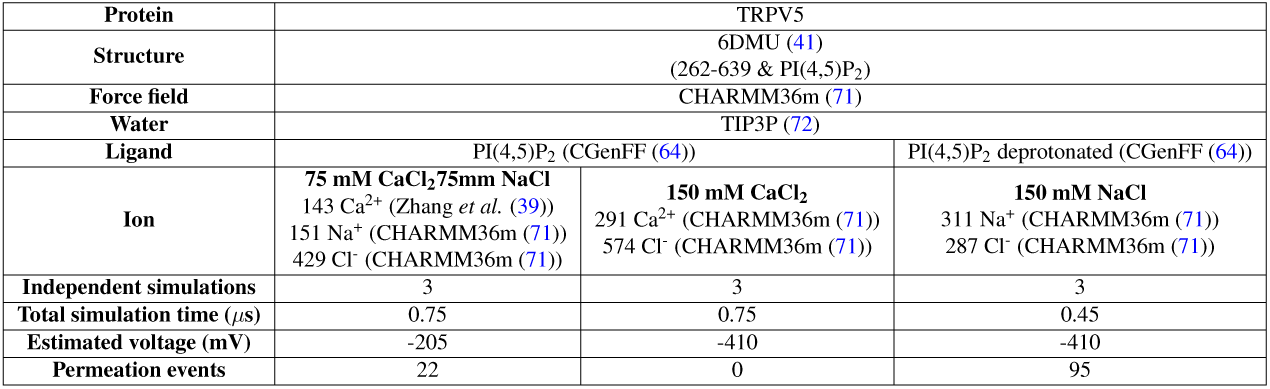
Summary of simulation details of additional control simulations of Ca^2+^-selective TRPV channels.

### Average permeation times for Ca^2+^ and Na^+^ cations in MD simulations of TRPV channels

The average permeation time of cations from mono-cationic simulations is described in Table S4. The average permeation time was defined as the time taken between the cation binding at binding site A, and dissociating from binding site C into the bulk solution. This data showed that for every TRPV system, the average permeation time was greater for Ca^2+^ permeation than for Na^+^, with the exception of TRPV6 where there was no discernible difference between the average permeation times. This greater permeation time of Ca^2+^ cations is likely a result of their greater affinity for the cation binding sites. However, the difference in permeation time is less than one would expect based on the residency times (Figure 3). For example, the average time for a Ca^2+^ cation to permeate through TRPV5 is ∼ 1.5-fold slower than Na^+^ permeation, whereas the residency time of cations at binding site A of TRPV5 would suggest a 12-fold difference. This supports the concept of co-operativity between successive binding sites increasing the unbinding rate of ions from binding sites, as proposed by Hille (7).

**Table S4.** Average time to permeate through the TRPV pore, as defined by the *z* position between binding sites A and C. The mean permeation time and standard error of the mean were calculated from five-fold replicated 250 ns simulations in mono-cationic 150 mM CaCl_2_ or 150 mM NaCl.

|  | Permeation time (ns) |  |
| --- | --- | --- |
|  | Ca<sup>2+</sup> | Na<sup>+</sup> |
| TRPV2 | 28.4 ± 7.6 | 7.0 ± 1.8 |
| TRPV3 | 6.3 ± 1.2 | 2.8 ± 0.2 |
| TRPV5 | 28.4 ± 3.9 | 18.2 ± 3.8 |
| TRPV6 | 12.0 ± 1.0 | 12.1 ± 1.5 |

### Pore-architecture of Ca^2+^-selective and non-selective TRPV channels from MD simulations

To investigate the time-averaged architecture of TRPV channels in our simulations, we used CHAP (47) to create profiles of the pore radius and the hydrophobicity of pore-facing residues (Figure S1). The CHAP analysis of the simulated TRPV structures showed similar profiles to the respective starting, static structures. These include: an upper constriction formed by the SF and a lower constriction formed by a hydrophobic gate.

**Fig. S1.**
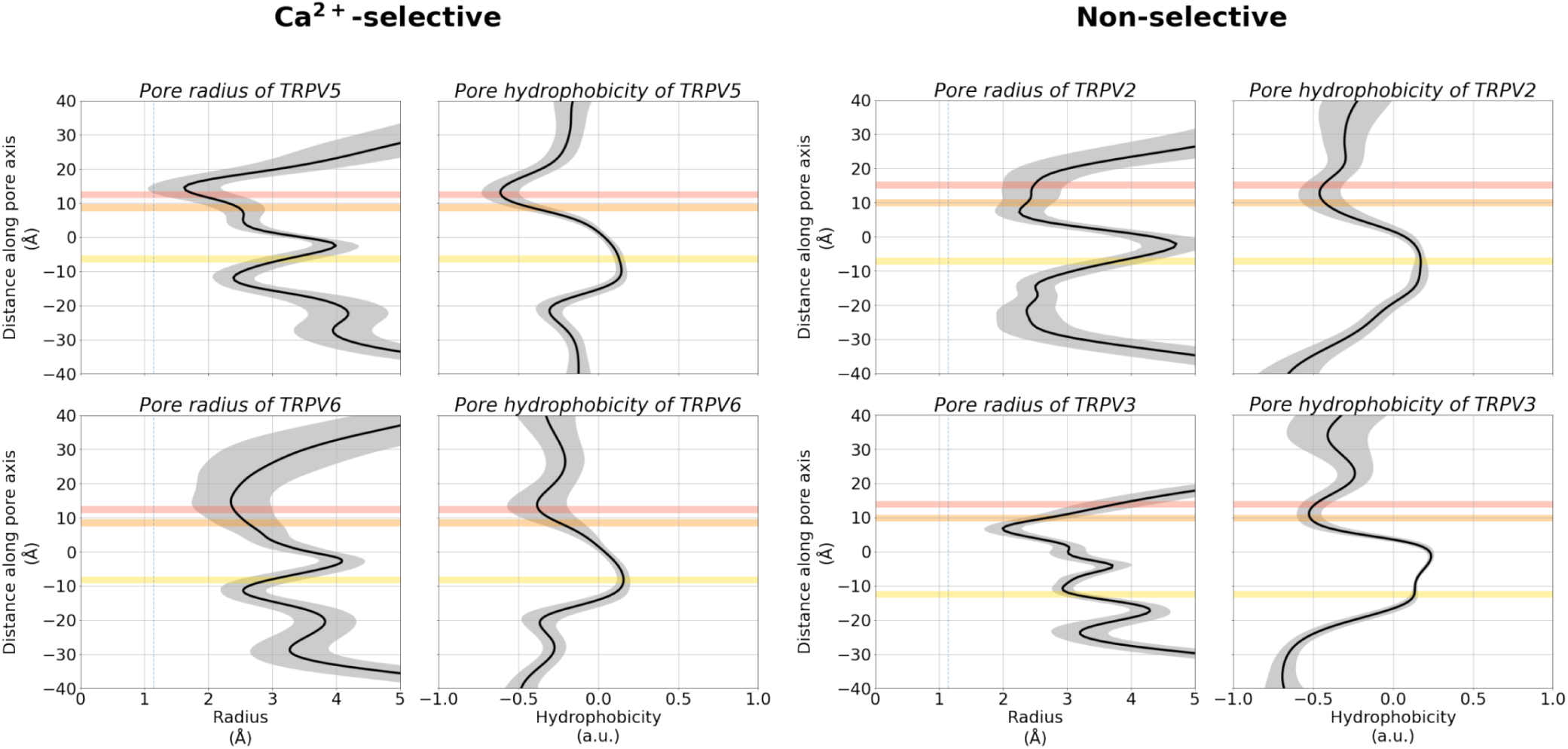
Pore architecture of TRPV channels from MD simulations, showing the average radius and hydrophobicity of the channel with respect to the relative *z* coordinate, obtained using CHAP (47). The mean radius or hydrophobicity (*black*) and standard deviation (*grey*) were calculated from concatenated trajectories of five-fold replicated 250 ns simulations in 150 mM CaCl_2_ with a 200 ps time-step. The shaded grey region represents the standard deviation. The average position of binding sites A, B, and C are shown as shaded red, orange, and yellow regions, respectively.

### Identification of cation binding sites using PENSA

In addition to permeation traces (Figure 2), cation binding sites were further identified using the *Featurizer* function of PENSA (44). This analysis identified cation binding sites from 3D density maxima in the simulation. As can be seen in Figure S2, this analysis identified binding sites A, B, and C, as well as cation binding sites within the extracellular loops of the protein. This finding is in agreement with previous suggestions of “recruitment sites” in TRPV6: negatively charged or polar residues that attract cations and funnel them towards the entrance of the pore (45, 46). In particular, our MD simulations suggested that the following residues act as recruitment sites: E522, D525, T528, F531, S532, E535, Y547, and Y549.

**Fig. S2.**
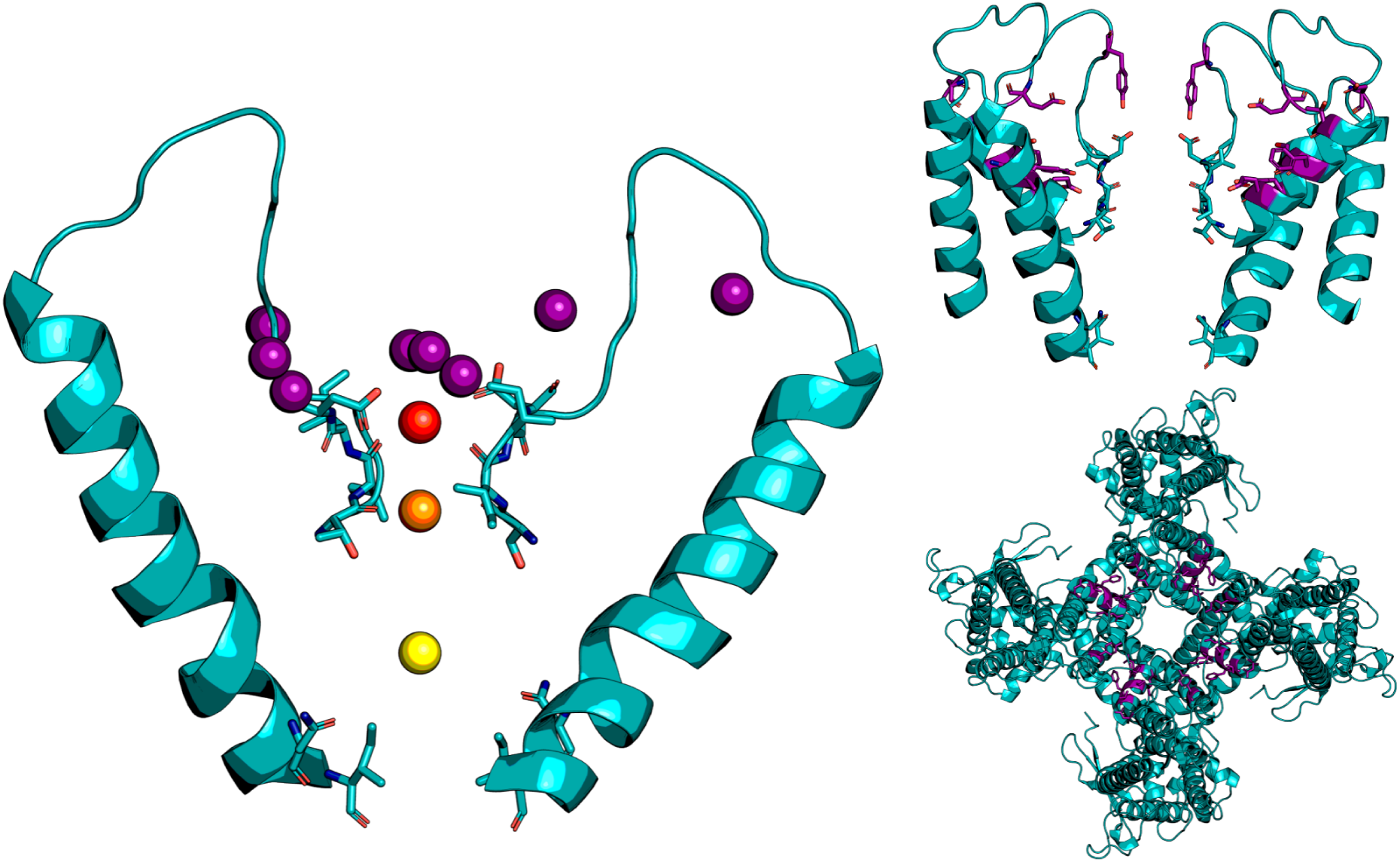
Cation binding sites in TRPV5 identified by PENSA. 3D density maxima of Ca^2+^ cations within 7 Å of the protein was analysed to identify 12 cation binding sites shown as pseudo-atoms (*left*). This analysis identified binding sites A (*red*), B (*orange*), and C (*yellow*) and several other “recruitment sites” (*purple*). The location of these recruitment sites (*purple*) help to attract cations and funnel them towards the pore entrance (*top right and top left*.

### W583 does not constitute a cation binding site in our simulations

In the structure of TRPV5, Hughes *et al*. reported a constriction below the hydrophobic lower gate formed by residue W583 (41). An analogous constriction formed by W583 can be observed in the structure of TRPV6 determined by McGoldrick *et al*. (42). Therefore, we investigated whether this constriction also constituted a cation binding site in our simulations (putatively referred to as binding site D), in addition to binding sites A, B, and C. Analysis of simulations of TRPV5 in a mono-cationic solution of 150 mM CaCl_2_ showed that the occupancy probability and residence time of Ca^2+^ at binding site D is substantially lower than at the other binding sites (Figure S3). Therefore, we did not consider W583 to constitute a functionally important cation binding site.

**Fig. S3.**
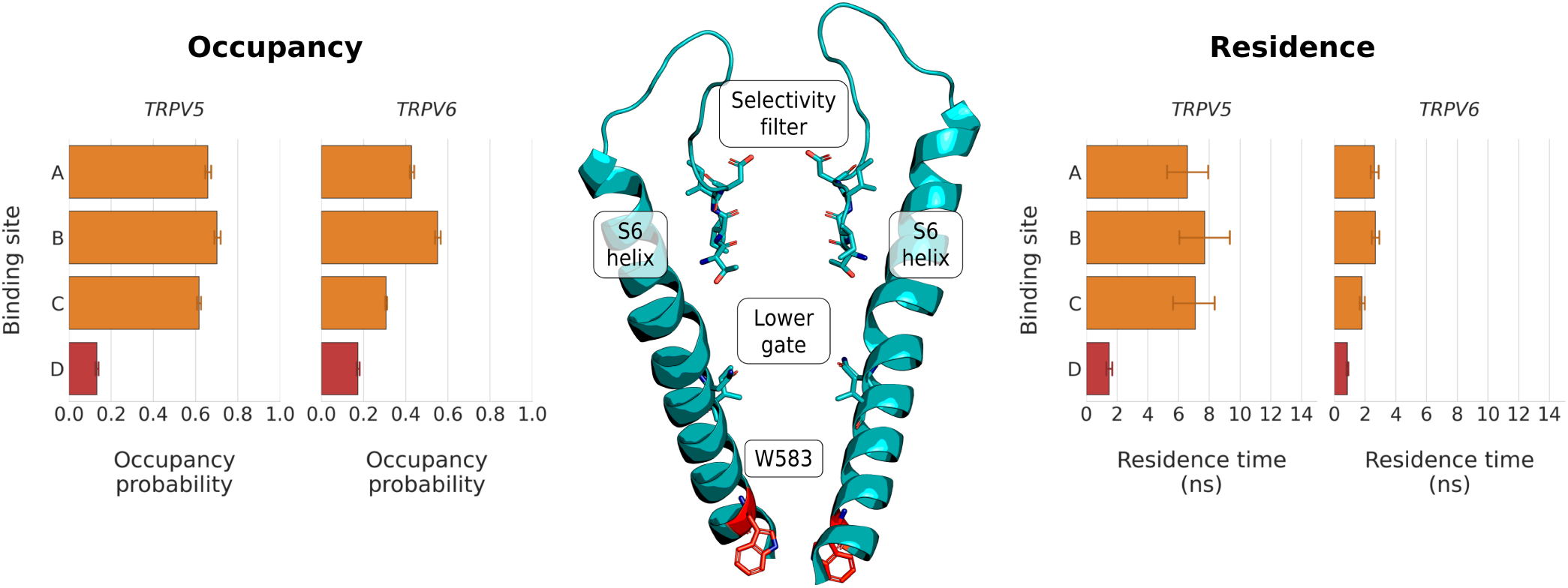
W583 does not form a functionally important cation binding site in simulations of TRPV5 in 150 mM CaCl_2_. W583 is located on the S6 helix, below the hydrophobic lower gate (*centre*). Analysis of the occupancy probability (*left*) and the residence time (*right*) showed that the constriction formed by W583 does not coordinate Ca^2+^ cations as efficiently as binding sites A, B, and C in our simulations.

### State-specific information and total correlation of cation binding site transitions in cation permeation

Table S5 shows all calculated excess SSI values and, in addition, normalised excess SSI for all channels and both ions investigated according to *exSSI*(*A, B*)_*norm*_ = *exSSI*(*A, B*)*/exSSI*(*A, B*)_*max*_.

**Table S5.** Calculated *exSSI* and *exSSInorm* values of cation transition from binding sites in MD simulations of TRPV channels. The mean *exSSI* or *exSSInorm*, and standard error of the mean, were calculated from five-fold replicated 250 ns simulations in mono-cationic solutions of 150 mM CaCl_2_ or 150 mM NaCl.

| Protein | Cation | $exSSI(A, B)$ | $exSSI(A, B)_{norm}$ | $exSSI(B, C)$ | $exSSI(B, C)_{norm}$ |
| --- | --- | --- | --- | --- | --- |
| TRPV2 | $Ca^{2+}$ | $0.15 \pm 0.06$ | $0.12 \pm 0.05$ | $0.88 \pm 0.13$ | $0.44 \pm 0.03$ |
| | $Na^+$ | $0.26 \pm 0.13$ | $0.18 \pm 0.06$ | $0.13 \pm 0.03$ | $0.18 \pm 0.02$ |
| TRPV3 | $Ca^{2+}$ | $0.34 \pm 0.09$ | $0.17 \pm 0.04$ | $0.96 \pm 0.17$ | $0.32 \pm 0.05$ |
| | $Na^+$ | $0.05 \pm 0.04$ | $0.03 \pm 0.02$ | $0.26 \pm 0.16$ | $0.10 \pm 0.06$ |
| TRPV5 | $Ca^{2+}$ | $1.25 \pm 0.15$ | $0.50 \pm 0.05$ | $1.01 \pm 0.25$ | $0.40 \pm 0.06$ |
| | $Na^+$ | $1.05 \pm 0.31$ | $0.30 \pm 0.06$ | $0.84 \pm 0.17$ | $0.28 \pm 0.04$ |
| TRPV6 | $Ca^{2+}$ | $0.81 \pm 0.16$ | $0.31 \pm 0.03$ | $0.53 \pm 0.25$ | $0.27 \pm 0.04$ |
| | $Na^+$ | $1.55 \pm 0.18$ | $0.39 \pm 0.03$ | $0.88 \pm 0.10$ | $0.32 \pm 0.03$ |

To evaluate and quantify the overall co-operativity in a system across all coupled binding site transition events, we used the concept of *total correlation*. This total correlation analysis showed that the Ca^2+^-selective TRPV channels had a greater total correlation than the non-selective TRPV channels (Figure S4). The greater total correlation in Ca^2+^-selective TRPV channels is due to these channels consisting of a three binding site knock-on permeation mechanism. However, non-selective TRPV channels consist of a two binding site knock-on permeation mechanism, due to the loss of co-operativity between binding sites A and B, reducing the total correlation of these channels.

**Fig. S4.**
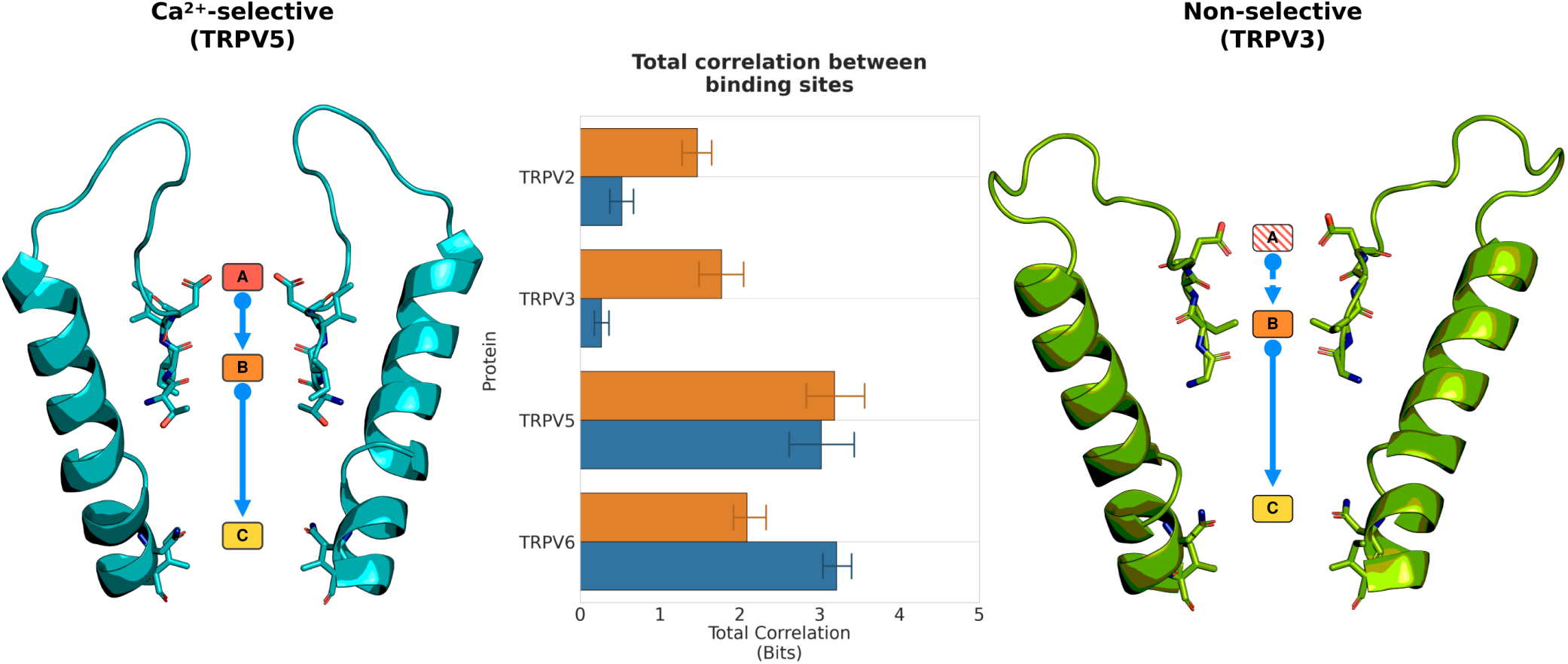
Total correlation of cation permeation between cation binding sites from simulations of TRPV channels. Comparison of the total correlation showed that the Ca^2+^-selective TRPV channels have greater total correlation than non-selective TRPV channels (*centre*). This greater total correlation in Ca^2+^-selective TRPV channels is a consequence of a knock-on mechanism between three binding sites (*left*). However, in non-selective TRPV channels, cation co-ordination at binding site A is reduced, resulting in reduced co-operativity, and a two binding site knock-on mechanism between binding sites B and C only (*right*).

### Lower voltage simulations of TRPV5 in a di-cationic solution

As described previously, TRPV5 and TRPV6 are highly Ca^2+^-selective with a reported *in vitro* experimental permeability ratio (*P*_*Ca*_*/P*_*Na*_) of ∼ 100:1 (20, 21). The remaining members of the TRPV subfamily, TRPV1-4, are still slightly Ca^2+^-selective however, with with a permeability ratio *P*_*Ca*_*/P*_*Na*_ of ∼ 10:1 (28). Table S6 shows the calculated *P*_*Ca*_*/P*_*Na*_ from our MD simulations of TRPV channels in di-cationic solutions at a voltage of -410 mV.

**Table S6.** Selectivity ratios of Ca^2+^ and Na^+^ permeation events from simulations of TRPV channels in a di-cationic solution.

| | $P_{\text{Ca}}/P_{\text{Na}}$ |
| --- | --- |
| **TRPV2** | 5.0 |
| **TRPV3** | 1.7 |
| **TRPV5** | 2.6 |
| **TRPV6** | 4.4 |

These calculated *in silico* selectivity values are lower than the *in vitro* selectivity values. We surmised that this may be due to the higher voltages (∼ 410 mV) used in our simulations to enhance the sampling rate of ion permeation events. To test this, we ran simulations of TRPV5 in a di-cationic solution using a reduced voltage of ∼ 205 mV. These simulations showed an increase in Ca^2+^ selectivity, as well as an increase in occupancy probability of Ca^2+^ cations in the cation binding sites (Figure S5). However, these simulations with a lower voltage resulted in a lower number of permeation events, meaning that they were not used to elucidate the mechanism of cation permeation due to reduced sampling.

**Fig. S5.**
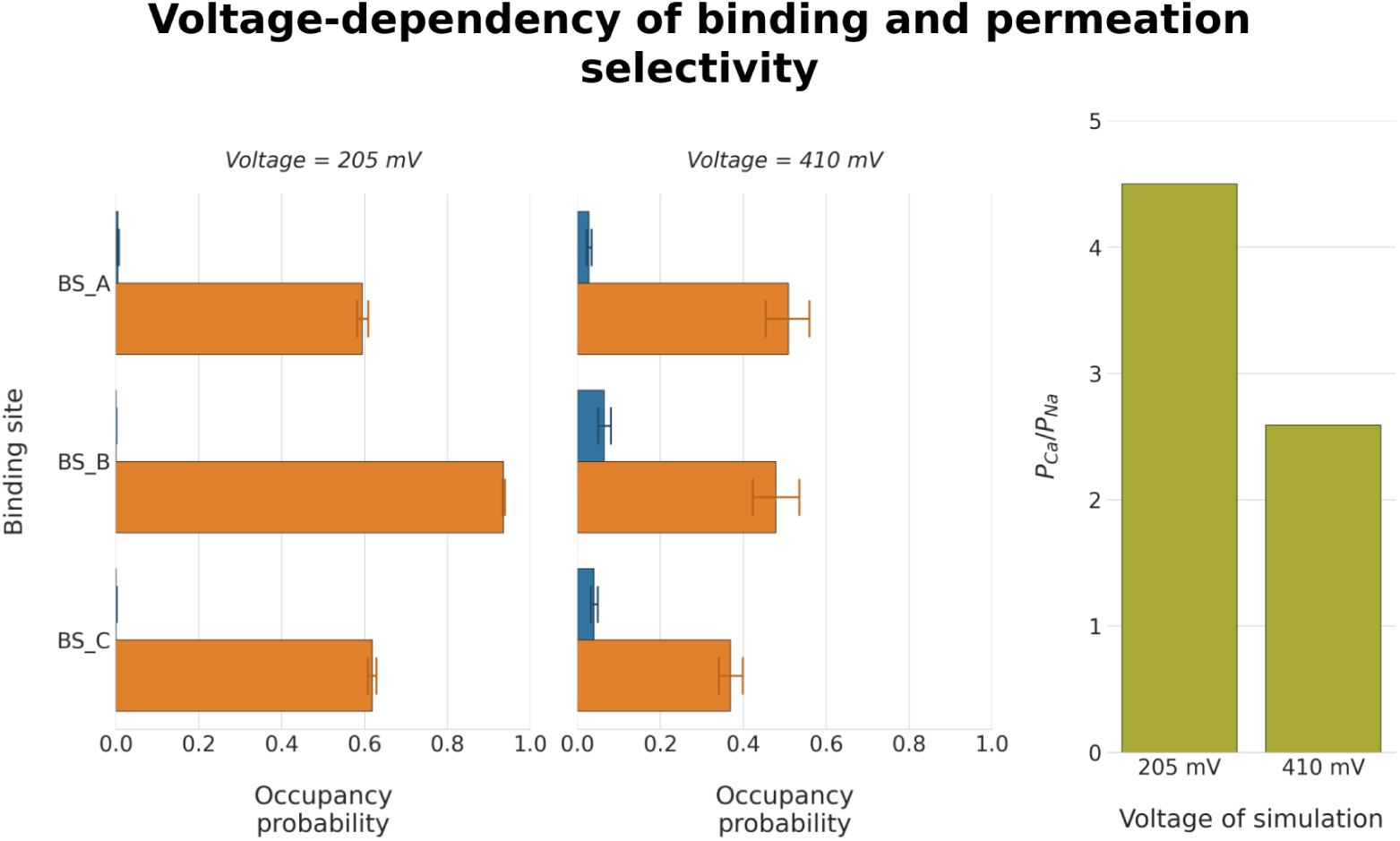
The effect of high voltage on Ca^2+^-selectivity in simulations of TRPV5 in di-cationic solutions. Simulations performed at a lower voltage of ∼ 205 mV (*left*) resulted in an increased occupancy probability of Ca^2+^ cations (*orange*) and a reduced occupancy probability of Na^+^ cations (*blue*) compared to simulations at a higher voltage of ∼ 410 mV (*centre*). This resulted in increased Ca^2+^-selectivity in lower voltage simulations, as summarised in the *P*_*Ca*_*/P*_*Na*_ value (*right*).

### Conformational flexibility in the selectivity filter of Ca^2+^-selective and non-selective TRPV channels

In addition to our investigations of the differences in SF architecture between Ca^2+^-selective and non-selective TRPV channels, we also investigated whether there were differences in the flexibility of the SFs. Analysis of the root mean square fluctuation (RMSF) of the backbone atoms for each residue in the SF showed similar RMSF values for all residues of all TRPV channels (Table S7), with RMSF values ranging between 0.6 Å and 1.3 Å.

**Table S7.** Root mean square fluctuation (RMSF) of the backbone of SF residues of TRPV channels from MD simulations. The mean RMSF and standard error of the mean for each residue was calculated from five-fold replicated 250 ns simulations of each channel in 150 mM CaCl_2_.

| Protein | RMSF (Å) |  |  |  |
| --- | --- | --- | --- | --- |
| | α-position | β-position | γ-position | δ-position |
| TRPV2 | 0.8 ± 0.04 | 0.7 ± 0.04 | 0.6 ± 0.05 | 0.7 ± 0.05 |
| TRPV3 | 0.9 ± 0.04 | 0.8 ± 0.05 | 0.7 ± 0.02 | 0.7 ± 0.02 |
| TRPV5 | 1.0 ± 0.07 | 0.7 ± 0.04 | 0.7 ± 0.03 | 0.7 ± 0.02 |
| TRPV6 | 1.3 ± 0.08 | 0.9 ± 0.04 | 0.9 ± 0.04 | 0.9 ± 0.03 |

In contrast, analysis of the backbone dihedral angles showed some differences in the angle distribution between Ca^2+^ and non-selective TRPV channels (Figure S6). The most obvious difference was in the *β*-position of the SF, where non-selective TRPV channels showed wider distribution of the *ϕ* angle, and principally on the *ψ* angle.

Comparison of the amino acid composition of the SF of TRPV channels shows that the *β*-position residue in Ca^2+^-selective TRPV channels is usually an isoleucine residue; however, in non-selective TRPV channels this *β*-residue is a much smaller glycine residue (Figure 6). As the sidechain of the *β*-position residue faces into hydrophobic pocket between the SF and pore-helix, the hydrophobic sidechain of isoleucine is involved in hydrophobic interactions with this region, increasing the backbone stability of the *β*-position in Ca^2+^-selective TRPV channels. On the other hand, the glycine residue at the *β*-position in the non-selective TRPV channels will have greater backbone flexibility. Furthermore, the increased flexibility at this *β*-position will affect the adjacent *α*- and *γ*-positions, explaining why these two positions show differences in their backbone dihedral angle distributions between Ca^2+^-selective and non-selective TRPV channels, but no difference in the *δ*-position.

**Fig. S6.**
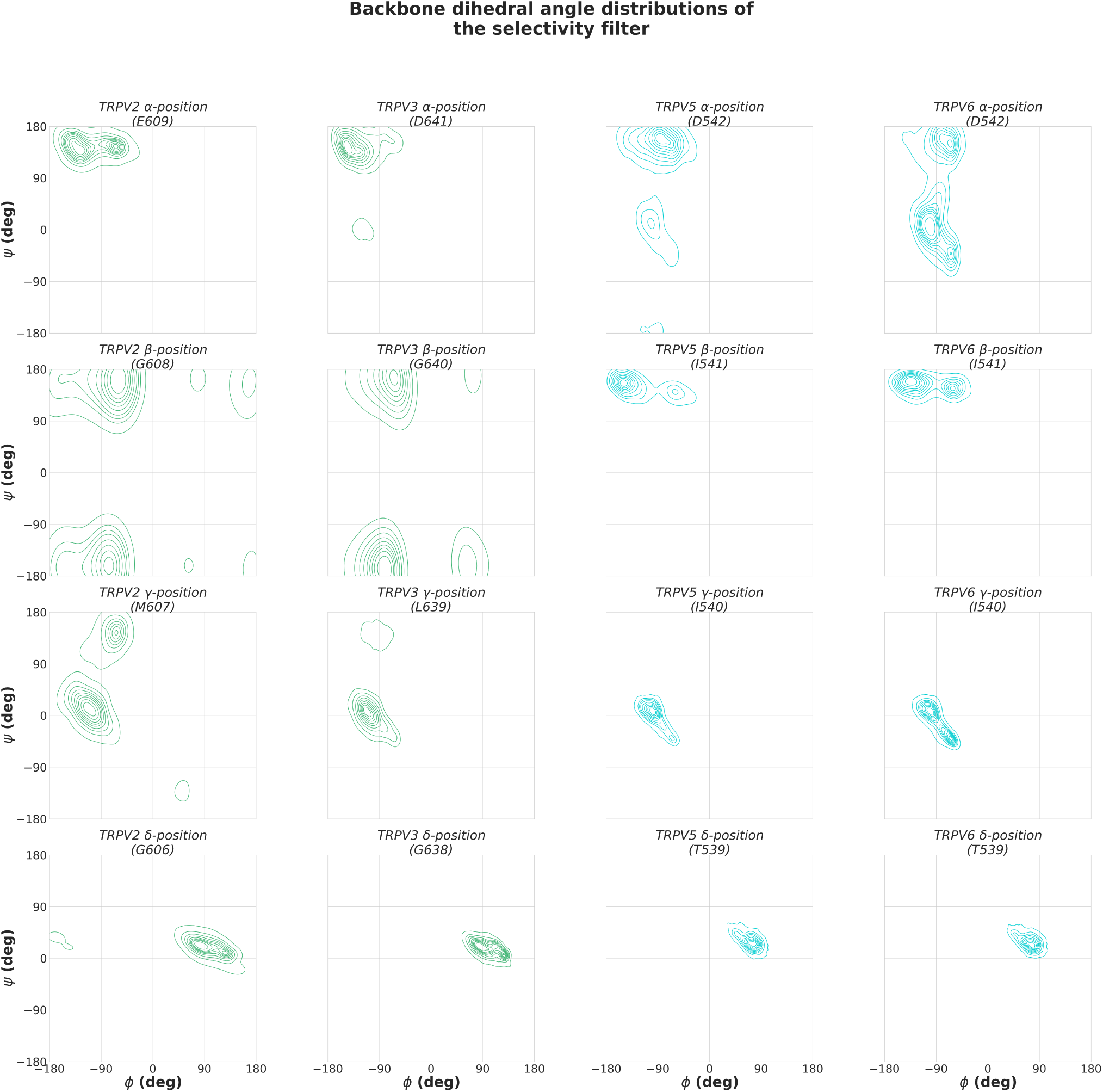
Backbone dihedral angle distribution of SF residues in TRPV channels. The *ϕ* and *ψ* angles were calculated across five-fold replicated 250 ns simulations of each TRPV channel in 150 mM CaCl_2_.

### The effect of PI(4,5)P_2_ protonation state on the TRPV5 simulation system

The cryo-EM structure utilised of TRPV5 of *Oryctolagus cuniculus* was solved to a resolution of 4 Å and resolved a PI(4,5)P_2_ molecule bound to each chain of the homotetrameric structure (41). At such a resolution, it is not possible to accurately model the protonation state of the PI(4,5)P_2_ molecule. Hughes *et al*. report that PI(4,5)P_2_ binding induces conformational changes related to channel activity. As our simulation protocol included harmonic restraints on the lower gate of the protein to maintain the open state of the structure, the protonation state of the PI(4,5)P_2_ would be inconsequential to the stability of the pore during our MD simulations. To confirm this, we performed three-fold 150 ns simulations of TRPV5 with deprotonated PI(4,5)P_2_ molecules, and compared the pore architecture to the production simulations of five-fold 250 ns simulations of TRPV5 with protonated PI(4,5)P_2_ molecules.

As can be seen in Figure S8, our simulations show no significant difference in the pore architecture dependent on the protonation state of the PI(4,5)P_2_ in TRPV5 simulations. This demonstrates that our simulation protocol and use of harmonic restraints on the lower gate are able to reliably constrain the protein structures in their open-state conformations.

**Fig. S7.**
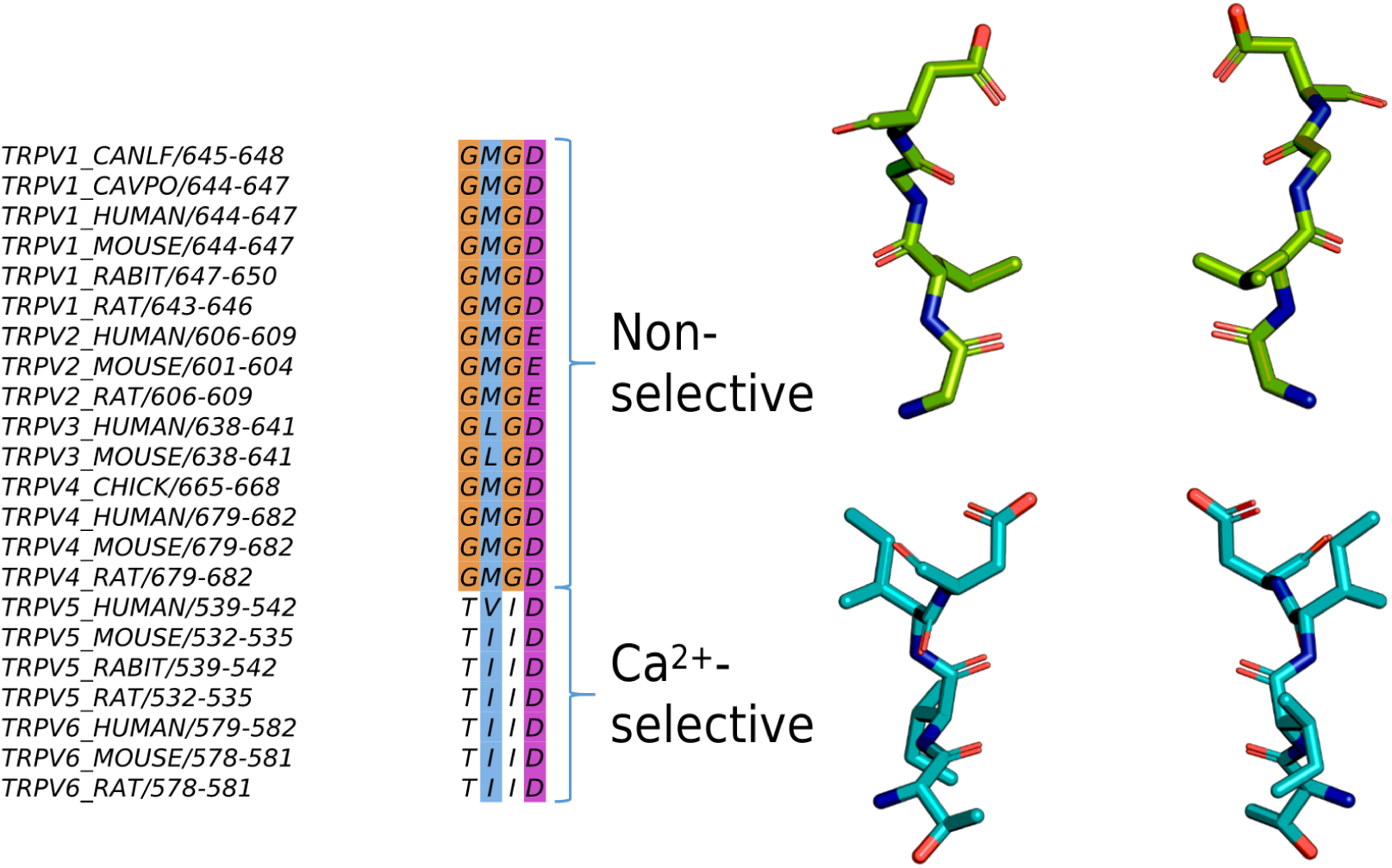
Multiple sequence alignment of the SF domains of TRPV sequences deposited in the Swiss-Prot database. Residues are coloured according to the ClustalX colouring scheme.

**Fig. S8.**
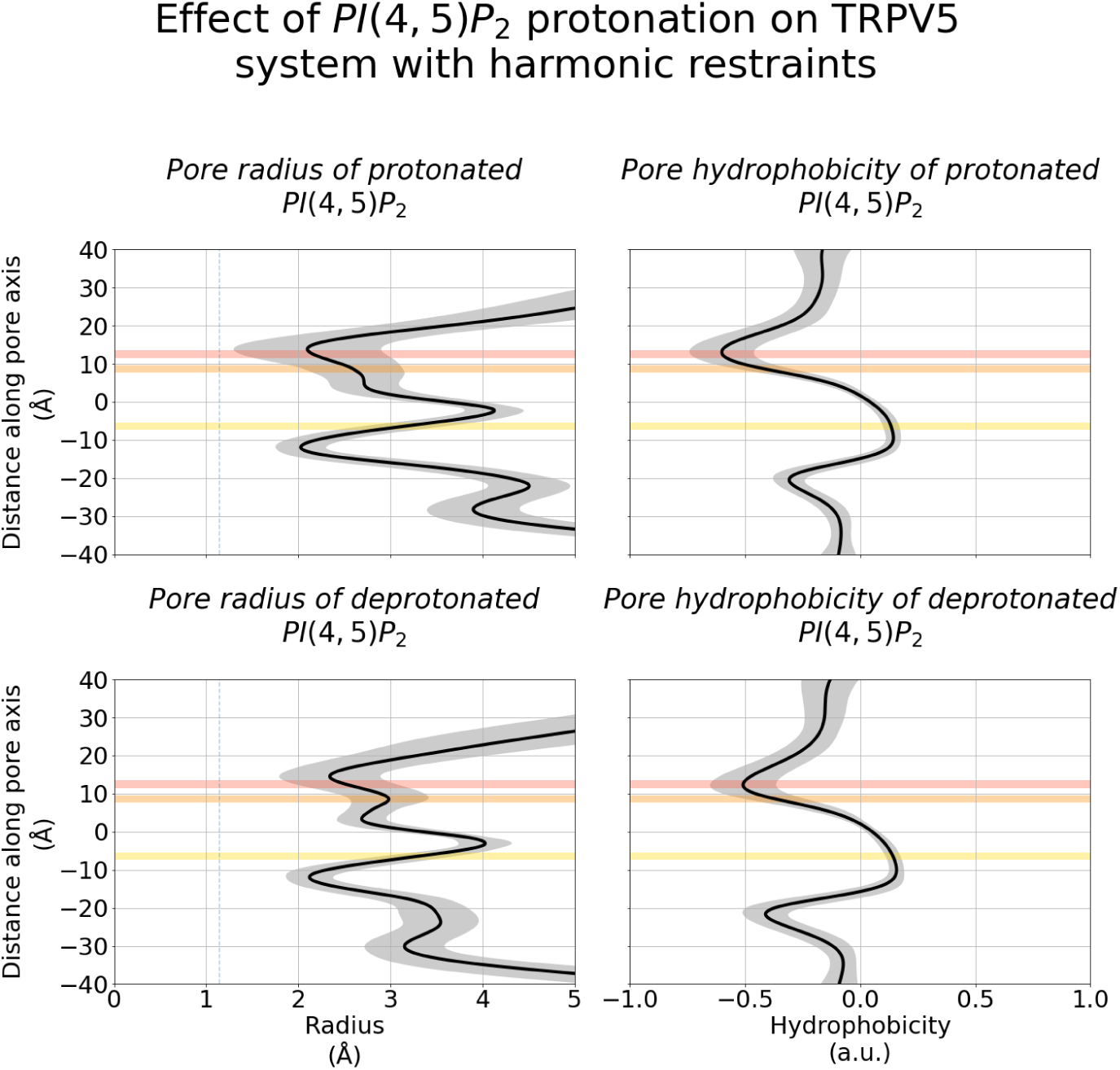
Pore architecture of TRPV5 channels from MD simulations with protonated (*top*) and deprotonated (*bottom*) PI(4,5)P2 molecules. The plots show the mean radius and hydrophobicity of the channel with respect to the relative *z* coordinate. The shaded grey region represents the standard deviation. The average position of binding sites A, B, and C are shown as shaded red, orange, and yellow regions, respectively.

